# Dopaminergic drugs modulate fear extinction related processes in humans, but effects are mild

**DOI:** 10.1101/2025.04.11.648372

**Authors:** Alice Doubliez, Kristina Köster, Lara Müntefering, Enzo Nio, Nicolas Diekmann, Andreas Thieme, Bilge Albayrak, Seyed Ali Nicksirat, Friedrich Erdlenbruch, Giorgi Batsikadze, Thomas Michael Ernst, Sen Cheng, Christian Josef Merz, Dagmar Timmann

**Affiliations:** Department of Neurology and Center for Translational Neuro- and Behavioral Sciences (C-TNBS); Department of Pediatrics I and C-TNBS, Essen University Hospital, University of Duisburg-Essen, Essen, Germany; Institute for Neural Computation, Ruhr University Bochum, Bochum, Germany; Department of Cognitive Psychology, Institute of Cognitive Neuroscience, Ruhr University Bochum, Bochum, Germany

**Keywords:** fear conditioning, dopamine, recall, memory consolidation, delay conditioning

## Abstract

The ability to extinguish learned fear responses is crucial for adaptive behavior. The mesolimbic dopaminergic system originating in the ventral tegmental area has been proposed to contribute to fear extinction learning because of its critical role in reward learning. The unexpected omission of aversive unconditioned stimuli (US) is considered as rewarding (outcome better than expected) and to drive extinction learning. We tested the hypothesis that extinction learning is facilitated by dopaminergic drugs and impeded by anti-dopaminergic drugs. The effects of dopamine agonists [levodopa (100 mg) and bromocriptine (1.25 mg)] and antagonists [tiapride (100 mg) and haloperidol (3 mg)] on fear extinction learning were compared to placebo in 146 young and healthy human participants. A three-day differential fear conditioning paradigm was performed with pupil size and skin conductance responses (SCRs) being recorded. Fear acquisition training was performed on day 1, extinction training on day 2, and recall was tested on day 3. The conditioned stimuli (CS+, CS-) consisted of two geometric figures. A short electrical stimulation was used as the aversive US. One of the four drugs or placebo was administered prior to the extinction phase on day 2. Overall, effects were small and seen only in the bromocriptine group. In line with our hypothesis, we measured reduced pupil dilation during late recall in the bromocriptine group compared to the placebo group, indicating faster re-extinction of spontaneously recovered fear reactions on the third day. Effects of levodopa and haloperidol were unspecific and related to generally increased SCR levels in the levodopa group (already prior to drug intake), and miotic side-effects of haloperidol. Findings provide additional support that the dopaminergic system contributes to extinction learning in humans, possibly by improving consolidation of fear extinction memory.

## Introduction

Learning to identify and react to threatening situations is essential for survival, but it is equally vital to adjust the behavioral responses when those stimuli are no longer associated with danger^1^. The inability to extinguish fearful memories is an inherent aspect of many anxiety disorders such as post-traumatic stress disorder and phobias^2,3^. A widely used experimental approach to study learned fear responses and their subsequent extinction is based on Pavlovian conditioning^4^. Fear conditioning assesses defensive responses elicited not only by the innate adverse stimulus itself but also by a formerly neutral stimulus predicting the occurrence of an adverse event. During fear acquisition training, an unconditioned stimulus (US), inducing innate fear, is repeatedly paired with a neutral conditioned stimulus (CS+), resulting in fear responses following CS onset. Fear extinction occurs when the CS+ is repeatedly presented in absence of the previously paired US, leading to a gradual decrease in learned fear conditioned responses (CRs). Fear conditioning can be studied in both animals and humans. Behavioral responses, such as freezing, are commonly assessed to measure fear learning and extinction learning in rodents. However, the strength of the US in human fear conditioning paradigms is rarely of sufficient magnitude to generate such defensive behaviors. Instead, physiological indicators like skin conductance responses (SCRs), heart rate, and pupillary responses, as well as subjective responses based on questionnaires are employed to assess fear-related learning^5^.

Classical theories of associative learning propose that new learning arises from the discrepancy between predicted and actual outcomes, corresponding to a prediction error (PE)^6^. It is well established that the mesolimbic dopaminergic pathways play a critical role in reward processing and reinforcement learning. The release of dopamine (DA), originating from the ventral tegmental area (VTA), in the nucleus accumbens (NAc) is a phenomenon that has been consistently observed towards unexpected rewarding stimuli in associative learning tasks^7–9^. In extinction learning, the absence of an expected aversive US results in a better-than-expected outcome, which may be internally perceived and processed as a reward PE^10–12^. Following this reasoning, it has been proposed that reward PE drive fear extinction learning and are also mediated by dopamine release from neurons originating in the VTA^10^. In support of this hypothesis, recent studies conducted in rodents reported activation of a subset of DA neurons in the VTA following unexpected US omission during fear extinction^13–16^, especially during early trials when the PE is the highest^13,15^. Moreover, optogenetically inhibiting or activating those DA neurons at the time of US omission is sufficient to impair or enhance, respectively, fear extinction learning^13^. Evidence suggests that the VTA DA neurons involved in fear extinction project and release DA in the NAc^17^. Further supporting this observation, direct administration of a D2 receptor antagonist, haloperidol, into NAc of rats was found to impair fear extinction^18^.

To understand the influence of DA on emotional learning processes in humans, dopaminergic pharmacological approaches during associative learning task have been employed^19–22^. These interventions typically involve the intake of DA agonists or antagonists that predominantly interact with either D1 or D2 receptors, thereby either enhancing dopaminergic effects (agonists) or inhibiting them (antagonists). Lissek and colleagues evaluated the systemic effect of the DA antagonist tiapride and DA agonist bromocriptine during a cognitive predictive learning task. Drugs were administered prior to extinction. Whereas tiapride intake impaired extinction learning when occurring in a novel context^20^, bromocriptine administration resulted in a significantly higher level of renewal in participants exhibiting renewal effects^21^ (i.e., return of the extinguished association in the acquisition context). These findings suggest that inhibiting DA leads to difficulties in extinction of previously learned associations when the context changes, while increasing DA level enhances the consolidation of memory associations. Likewise, in a fear conditioning paradigm, administration of levodopa following fear extinction training and thus targeting extinction memory consolidation resulted in an enhancement of extinction memories during recall in humans^19,22^. Findings appeared to depend on the success of fear extinction learning^23^.

To further investigate the effect of the dopaminergic system on fear extinction learning, we conducted a three-day fear conditioning study in healthy human participants who received a systemic single dose of either a DA agonist, a DA antagonist or a placebo prior to fear extinction training. Consistent with the known dopaminergic signals in reward-associated learning, we hypothesized that DA agonists enhance fear extinction learning and thus reduce recall of learned fear responses, whereas DA antagonists impair fear extinction learning and increase recall of learned fear responses following extinction learning.

## Materials and methods

### Participants

160 young and healthy participants (18–35 years) were recruited on university campuses and through online and offline advertisements. They received monetary compensation for their participation. Four participants were excluded due to technical errors, five for not completing all three days of the study, one for failing to show contingency awareness after fear acquisition training, and two due to high scores on the Depression Anxiety Stress Scale (DASS-21-G)^24^. A total of 146 participants (74 women and 72 men, *Table 1*) were included in the final analysis. All participants were fluent in German, non-smokers, reported no intake of medication or illicit drugs affecting the central nervous system, had no personal or family history of neurological or psychiatric disorders, had no contraindications to dopaminergic/anti-dopaminergic drugs, and had never taken part in similar learning experiments. A physician evaluation and electrocardiogram, including QTc interval assessment, were conducted to rule out cardiac contraindications. Additionally, female participants were required not to be pregnant, breastfeeding, or using hormonal contraceptives. On day 1 participants completed the DASS-21- G scale; those who scored above moderate threshold (>20 for depression, >14 for anxiety and/or >25 for stress components) were excluded. The study was approved by the University Hospital Essen ethics committee and was conducted in accordance with the Declaration of Helsinki. Informed consent of participants was obtained by a physician before the experiment.

**Table 1.** Demographic characteristics and Depression Anxiety Stress Scale Scores (DASS-21-G) across medication groups. Group characteristics are presented as mean (± standard deviation), unless otherwise specified. The education level indicates the total time spent in the education system. The Edinburgh Handedness Inventory details the number of participants classified as right-handed, left-handed, or ambidextrous (ambidext.).

|  | GROUP A |  |  | GROUP B |  |  |
| --- | --- | --- | --- | --- | --- | --- |
|  | Levodopa | Placebo A | Tiapride | Bromocriptine | Placebo B | Haloperidol |
| **Demographic** |  |  |  |  |  |  |
| Participants (n) | 24 | 25 | 22 | 25 | 25 | 25 |
| Female (%) | 54.2% | 56% | 50% | 48% | 44% | 52% |
| Age (years) | 24.54 ± 3.99 | 24.68 ± 4.36 | 23.68 ± 3.99 | 24.16 ± 3.42 | 24.92 ± 4.51 | 25.00 ± 3.65 |
| Education Level (years) | 15.42 ± 2.04 | 16.94 ± 3.62 | 15.70 ± 1.86 | 15.40 ± 2.67 | 16.74 ± 3.12 | 16.34 ± 3.10 |
| Edinburgh Handedness Inventory (n) | Right: 21<br>Left: 2<br>Ambidext.: 1 | Right: 22<br>Left: 2<br>Ambidext.: 1 | Right: 18<br>Left: 4<br>Ambidext.: 0 | Right: 23<br>Left: 1<br>Ambidext.: 1 | Right: 23<br>Left: 1<br>Ambidext.: 1 | Right: 21<br>Left: 4<br>Ambidext.: 0 |
| **Depression Anxiety Stress Scale** |  |  |  |  |  |  |
| Depression | 3.54 ± 2.25 | 2.6 ± 1.76 | 3.09 ± 2.35 | 3.40 ± 2.66 | 3.16 ± 2.53 | 3.04 ± 2.46 |
| Anxiety | 2.92 ± 1.91 | 2.16 ± 1.37 | 2.77 ± 1.66 | 2.04 ± 1.06 | 2.20 ± 1.47 | 2.16 ± 1.68 |
| Stress | 4.46 ± 2.93 | 4.04 ± 2.26 | 4.00 ± 3.02 | 4 ± 2.60 | 3.96 ± 2.70 | 4.64 ± 3.15 |

### Study design

All participants completed the same three-day differential fear conditioning paradigm while their pupil size and skin conductance responses (SCRs) were recorded (*Figure 1–A*). As the times to peak concentration differed between the four drugs, participants were randomly assigned to two groups (A and B; *Table 1*). While group A participants received either placebo, levodopa or tiapride, group B participants received either placebo, bromocriptine or haloperidol. Participants were randomly assigned to one of three drug groups, with randomization ensuring balanced sex distribution across groups. Group A performed the experiment under low lighting conditions, whereas Group B performed the experiment under ambient lighting conditions.

**Figure 1.**
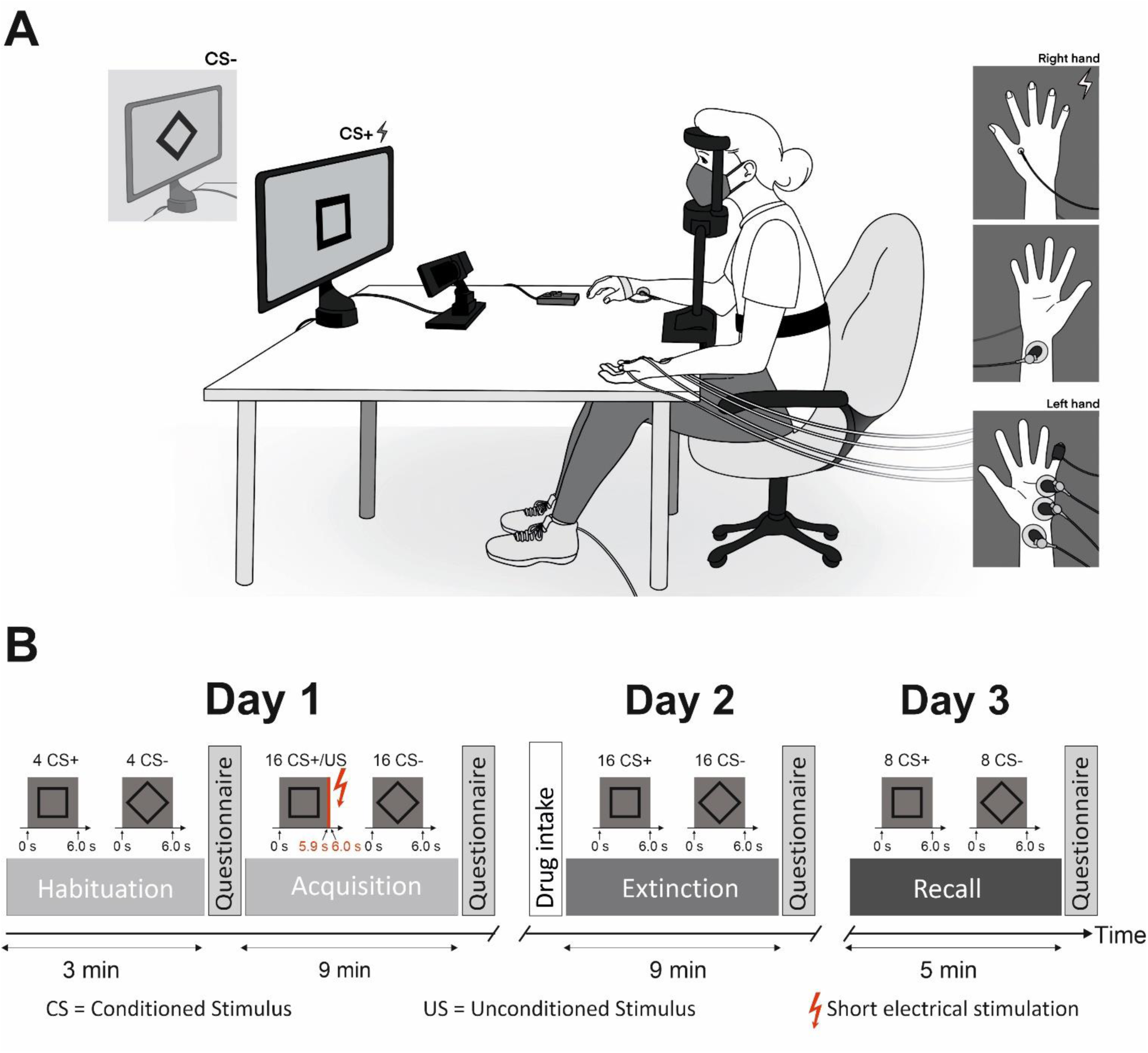
(A) **Illustration of the experimental setup** representing participants sitting in front of a computer screen displaying black-and-white geometric figure trials while their pupil size and skin conductance (SCR) were recorded. Note that participants wore face masks because the experiment was performed during the COVID19 pandemic. (B) **Three-day fear conditioning paradigm.** On day 1, participants underwent habituation and acquisition training. Day 2 begins with drug intake and is followed by extinction training. On day 3, the recall phase takes place.

Pupil size was monitored using an eye tracking system (EyeLink® 1000 Plus, SR Research Ltd., Ontario, Canada), positioned 60 cm from participants’ eyes. Participants used a headrest to maintain alignment with the camera and display screen. Additionally, calibration of the eye tracking system was conducted prior each phase. Skin conductance was recorded through an MP160 Data Acquisition Hardware unit (BIOPAC Systems Inc, Goleta, CA), with two electrodes affixed to the participant’s left-hand hypothenar eminence (*Figure 1-A*).

A short electrical stimulation, delivered as the aversive unconditioned stimuli (US), was generated by a constant current stimulator (DS7A, Digitimer Ltd., London, UK) and applied to the right hand’s first dorsal interosseous muscle via a 6.5 mm concentric surface electrode (WASP electrode, Specialty Developments, Bexley, UK). The electrode position was marked on day 1 for consistent positioning on following days. The 100 ms US consisted of four consecutive 500 µs pulses (maximum output voltage: 400 V) with 33 ms interval. Before the experiment, the stimulation intensity was adjusted individually to be highly uncomfortable but not painful. US intensity was increased by 20%, same as in Inoue et al.^25^, and remained the same for the three days of the experiment. The electrical stimulation electrode was attached throughout the experiment providing stimulation only during paired CS+/US trial. Following local rules, participants and investigators wore face masks and had a daily negative COVID-19 rapid test. This study was preregistered on the Open Science Framework (OSF) (https://doi.org/10.17605/OSF.IO/A3KJN).

The sample size required was determined for each group (A and B) separately using G*Power 3.1.9.7 for repeated measures analyses of variance (ANOVA) within-between interactions. Our goal was to obtain 0.85 power (1 − β) to detect a medium effect size f = 0.25^26,27^ at the standard 0.05 alpha error probability and an assumed correlation among repeated measurements of r = 0.20^28^.

### Fear conditioning paradigm

This differential fear conditioning paradigm was adapted from Ernst et al.^29^. Throughout the three-day fear conditioning, participants were instructed to keep their heads still and focus on the computer screen displaying the paradigm using Presentation software (version 22.1, Neurobehavioral System Inc, Berkeley, CA). Two pictures of black-and-grey geometric figures, a square and a diamond (square titled by 45°), were used as visual conditioned stimuli (either CS+ or CS-) (*Figure 1–A and B*). The visual CS paired with the electrical US was randomly assigned and remained the same throughout the experiment.

During the experiment, three types of trials were presented to the participants: CS+ followed by a US (paired CS+/US trial), CS+ without a US (CS+ only trial), and CS- that was never followed by a US. For CS+/US paired trials, the US was administered 5.9 s after CS+ onset. During the inter-trial interval (ITI), a black cross on a grey background was presented to the participants (ITI duration varied from 6 s to 12 s). The paradigm included a habituation phase (4 CS+ trials, 4 CS- trials, presented in alternating order) followed by fear acquisition training (16 paired CS+/US trials - 100% reinforcement rate, 16 CS- trials) on day 1, extinction training (16 CS+ trials, 16 CS- trials) on day 2, and a recall test (8 CS+, 8 CS-) on day 3 (*Figure 1-B*). During fear acquisition training, full reinforcement rate was used to allow for maximal prediction error in initial extinction training trials. The different trial types in each phase were presented in pseudo-randomized order, similar to Ernst et al.^29^.

### Drug treatment

Prior to extinction training on day 2, a single dose of levodopa/carbidopa (100/25 mg; referred to as “levodopa” hereafter)^30^, bromocriptine (1.25 mg)^21,31–34^, tiapride (100 mg)^20^, haloperidol (3 mg) or placebo was given to participants in non-transparent capsules. Drugs were administered at specific time prior extinction training assuring maximum serum levels at the beginning of extinction training (60 min for levodopa^35^; 90 min for bromocriptine^36^; 120 min for haloperidol^37^ and tiapride^38^). Prior to levodopa or bromocriptine intake, 20 mg domperidone^39^ was given to prevent nausea. To ensure a double-blind experiment, a placebo capsule was ingested at a corresponding time after tiapride or haloperidol intake, matching the total number of capsules ingested (*Table S1*). Participants fasted for 2 hours before drug intake, and blood samples were collected daily after experiment for external drug concentration analysis (*Table S2*).

### Questionnaires and contingency awareness

Participants were informed that electrical stimuli (US) may be administered, and that the CS and US presentations patterns would remain consistent throughout all phases. However, they were not informed about the CS/US contingencies or whether and when the US would be presented.

To evaluate contingency awareness, participants specified which of the two CSs had been followed by a US at the end of each phase where they reported receiving electric stimuli. Moreover, participants had to indicate whether a US was expected after the CS presentation and, if so, to estimate the number and percentage of US that occurred after the respective CS presentation (US expectancy) during that phase.

Regardless of US perception, participants used nine-step Likert scales to rate both CSs respective valence, arousal, fear, US expectancy and US unpleasantness at the end of all phases^40^.

### Skin conductance response analysis

To eliminate low-frequency drifts, skin conductance data were low-pass filtered at 10 Hz directly by the recording hardware system (EDA 100C-MRI, BIOPAC Systems Inc, Goleta, CA). Semi-automated peak detection was performed in MATLAB, and SCRs were defined as the maximum through-to-peak amplitude within a given time interval after CS onset^41^. A minimum amplitude criterion of 0.01 μS and a minimum rise time of 500 ms was used as the SCR detection threshold ^42^. Trials that did not meet the criteria were scored as zero and included in the subsequent data analysis. SCR was evaluated within a time window from 1.0 s to 5.9 s after CS onset. To account for between-subjects variance, the resulting raw SCR amplitudes were increased by 1 μS and normalized through a logarithmic transformation (LN(1+SCR))^42,43^.

### Pupillometry analysis

Preprocessing of the raw pupil data was performed in MATLAB (version 9.13 (R2022b), MathWorks, Natick, USA) in accordance with Kret and Sjak-Shie^44^ guidelines method. Only trials where eyes were opened and looking at the visual stimuli (CS) in the center of the screen were retained. Invalid samples were identified following multiple steps: 1. dilatation speed outliers and edge artifacts, which compares the change in pupil width relative to adjacent samples; 2. trendline deviation outliers, which observes the ratio of pupil area to adjacent samples; and 3. temporally isolated samples outliers, underlying single data points separated from adjacent samples. Interpolation of the missing data points was performed for gaps duration of less than 250 ms. Trials with major artifacts or many blinks were excluded from the analyses. Furthermore, a simultaneous visual examination of the raw and processed pupil data was carried out to ensure processing coherence and determine whether both eyes were properly recorded and should be kept in the following analysis. If one eye showed significantly more artifacts, it was omitted from the analysis. We focused our analysis on the 2 s preceding US onset, which correspond to the time interval that measures the largest difference between CS+ and CS- during fear acquisition training^28^. For each trial, the baseline was computed as the mean pupil size recorded during the 300 ms before CS onset and subtracted from the corresponding pupil size during CS^28,45^.

### Statistical analysis

Primary outcome measures included SCRs and pupil size. For each phase of the experiment—fear acquisition training, extinction training, and recall test—events of each kind (CS+ and CS-) were grouped into four-trials blocks representing the first (early) and the second (late) half of those phases (e.g. the 4 first CS+ trials of fear acquisition training correspond to early acquisition and the 4 last CS+ trials of fear acquisition training correspond to late acquisition). SCRs and pupil size were analyzed separately using non-parametric repeated-measures ANOVA-type statistics (ATS) via the PROC Mixed procedure in SAS (SAS Studio 3.8, SAS Institute Inc, Cary, NC, USA). These analyses included medication subgroup as a between-subjects factor and stimulus type (CS+ and CS-) and block (early and late phase) as within-subjects factors. Additionally, a separate non-parametric ATS analysis was conducted on the first trial of the recall phase, with medication subgroup as a between-subjects factor and stimulus type (CS+ and CS-) as a within-subjects factor.

Self-reported ratings were analyzed using non-parametric ATS for repeated measures. The rating type (valence, arousal, fear, or US expectancy) was treated as the dependent variable, with stimulus type (CS+ and CS−) and phase (post-habituation, post-acquisition, post-extinction, and post-recall) as within-subject factors, and medication group as a between-subjects factor.

To assess the effect of drug intake on baseline pupil size, we conducted a non-parametric ATS within each medication group to compare baseline pupil size across all experimental phases.

Significant results from the non-parametric ATS were followed by post-hoc comparisons using least square means tests. Dunnett’s adjustment was applied to control for multiple comparisons when comparing each treatment group to their respective control conditions. Tukey’s test was used when comparing other conditions within each medication group.

## Results group A

### Questionnaires

Across all participants, CS+ was rated significantly higher in unpleasantness, arousal, and fear compared to CS- after fear acquisition training, with these differences persisting through extinction and recall phases (*Figure S1*;*Table S3; Table S4*). Participants estimated a high likelihood of US occurrence following CS+ (98.21 ± 11.40%), while the probability was minimal for CS- (1.54 ± 5.34%; *Table S5*).

### Pupillometry

#### Habituation phase

During the habituation phase, the mean (differential) pupil size towards CS+ and CS- did not differ between groups (*Figure 2–A)*. Non-parametric ATS revealed no significant main effects of Stimulus (p*=*0.233), Group (p = 0.383), or Stimulus x Group (p = 0.613) interactions (Table S6).

**Figure 2.**
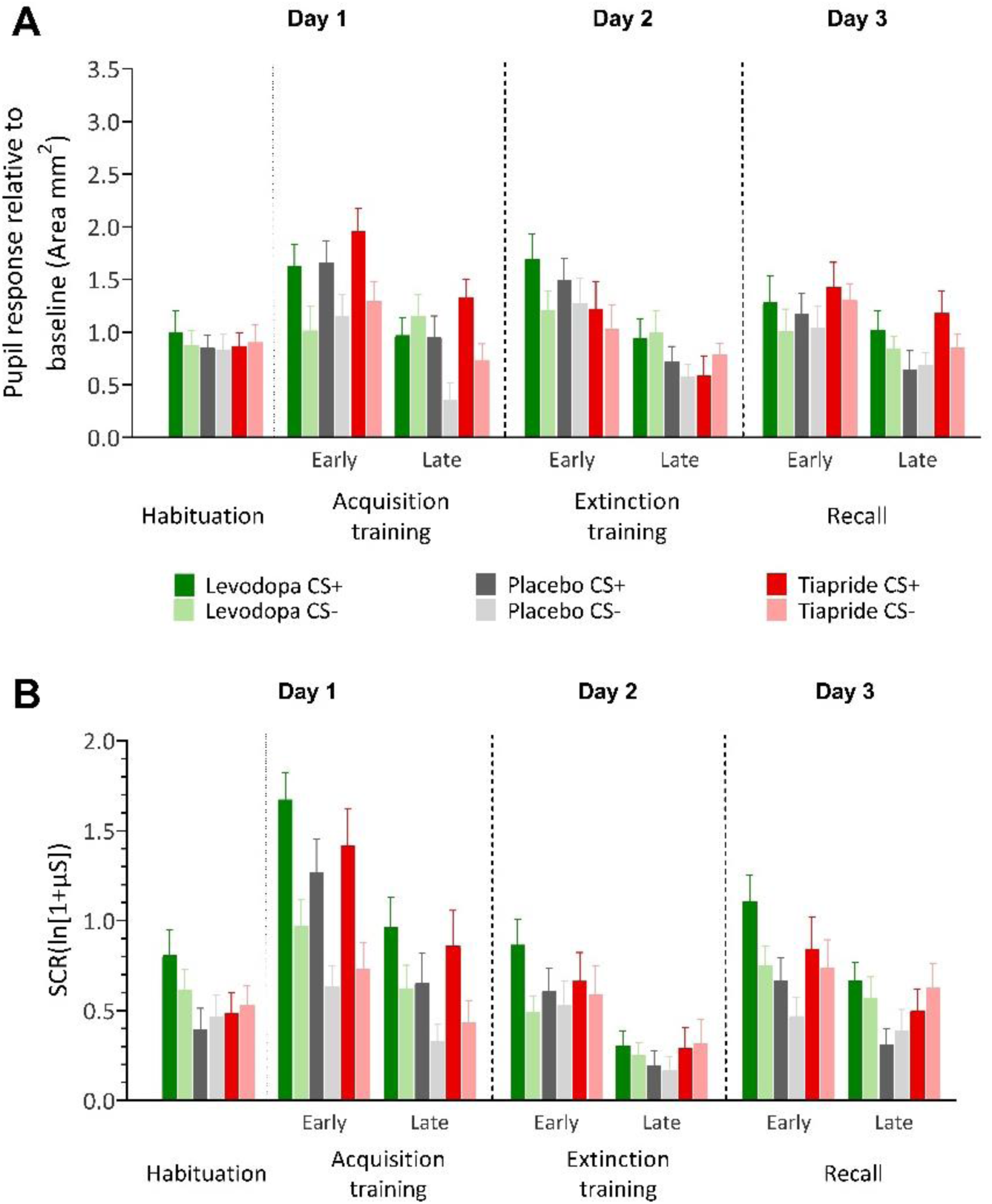
(A) **Pupil response** relative to baseline for habituation and early and late blocks of acquisition and extinction training as well as the recall phase in the groups receiving levodopa, placebo or tiapride. Bars represent means, error bars indicate S.E.M. (B) **Skin Conductance Responses** (SCRs) during habituation and early and late blocks of acquisition and extinction training as well as the recall phase in the groups receiving levodopa, placebo or tiapride. Bars represent means, error bars indicate S.E.M.

#### Fear acquisition training

During fear acquisition training, levodopa, placebo and tiapride groups showed significantly higher pupil responses to CS+ compared to CS- and during the early compared to the late part of the phase (*Figure 2-A*). Non-parametric ATS revealed a significant main effect of Stimulus (F_(1,55.5)_=37.40, p< 0.001) and Block (F_(1,66.7)_=32.25, p< 0.001), but no Group difference (p=0.300). Both Stimulus x Group (F_(1.86,65.1)_=3.22, p< 0.044) and Stimulus x Group x Block (F_(1.96,65.1)_=3.30, p < 0.038) interactions were significant, while other interactions (all p>0.050) were not statistically significant (*Table S6*). Stimulus x Block x Group post hoc analysis demonstrated significantly higher pupil responses during early CS+ compared to late CS+ (p<0.030), as well as in early CS+ compared to early CS- (p<0.012) in the placebo group. The levodopa group exhibited significantly higher pupil responses following late CS- compared to the placebo group for late CS- (p=0.010), with no difference observed for the CS+ stimuli. In comparison to the placebo group, no significant differences were observed for tiapride (all p>0.530) regarding both CS+ and CS- pupil responses during early or late phase. Note, however, that for SCRs no significant group interaction effects were found in fear acquisition training.

#### Extinction training

During extinction training, levodopa, placebo and tiapride groups showed significantly higher pupil responses during the early compared to the late part of the phase and in particular towards the CS+ compared to the CS- (*Figure 2-A*). Non-parametric ATS revealed a significant main effect of Block (F_(1,61.3)_=33.80, p<0.001), but no Stimuli (p=0.365) or Group difference (p=0.060; Table S6). Stimulus x Block (F_(1,63.1)_=5.50, p = 0.019) interaction demonstrated significantly higher CS+ in early compared to late blocks (all p<0.001), other interactions (all p>0.300) were not significant.

#### Recall

During recall, all three groups demonstrated significantly higher pupil size response during the early compared to the late part of the phase (*Figure 2-A*). There was no significant difference comparing groups or stimulus types. Non-parametric ATS revealed a main effect of early compared to late blocks (F_(1,52.2)_=15.19, p<0.001), while no main effects were observed for Stimulus type (p=0.330) and Group (p=0.130; *Table S6*). Additionally, none of the interactions were significant [Stimulus x Block (p=0.950), Group x Stimulus (p=0.879), Group × Block (p=0.473) or Group x Stimulus x Block (p=0.562)]. Likewise, reanalysis considering the first trial of recall only revealed no significant group differences (*Figure S2-A; Table S7*).

### Skin conductance responses (SCRs)

#### Habituation phase

During the habituation phase, the mean SCR amplitudes towards CS+ was higher in levodopa compared to placebo groups (*Figure 2–B*). Non-parametric ATS revealed no significant main effects of Stimulus (p=0.490) and Group (p=0.076), but a significant effect of Stimulus x Group (p=0.002) interaction *(Table S6*). Post hoc analysis of Stimulus x Group demonstrated no significantly higher SCR amplitude for both CS+ and CS- between all three groups (all p>0.088), except between levodopa CS+ and Placebo CS+ (p=0.012) of this phase.

#### Fear acquisition training

During fear acquisition training, levodopa, placebo and tiapride groups showed significantly higher mean SCR amplitudes to CS+ compared to CS- and during the early compared to the late part of the phase (*Figure 2–B*). Non-parametric ATS revealed a significant main effect of Stimulus (F_(1,67)_=65.10, p<0.001) and Block (F_(1,62.1)_=59.45, p<0.001), but no Group difference (p=0.066). Stimulus x Block (F_(1,65.1)_=6.17, p=0.013) interaction was significant, other interactions (all p>0.833) were not significant *(Table S6*). Post hoc analysis of Stimulus x Block demonstrated significantly higher CS+ compared to CS- SCR amplitude in both early and late (all p<0.001) as well as higher CS+ and CS- SCR amplitude in early compared to late blocks (all p<0.001).

#### Extinction training

During extinction training, levodopa, placebo and tiapride groups showed significantly higher mean SCR amplitudes towards the CS+ compared to the CS- and during the early compared to the late part of the phase (*Figure 2–B*). Non-parametric ATS revealed a significant main effect of Stimulus (F_(1,58.6)_=11.08, p<0.001) and Block (F_(1,60)_=65.60, p<0.001), but no Group difference (p=0.158). Stimulus x Block (F_(1,63.1)_=4.75, p=0.029) interaction showed significant differences, other interactions (all p>0.553) were not significant *(Table S6*). Post-hoc analysis of Stimulus x Block demonstrated significantly higher CS+ compared to CS- SCR amplitude in early block only (p<0.001; late block p=0.711) as well as higher CS+ and CS- SCR amplitude in early compared to late blocks (all p<0.001).

#### Recall

During recall, all three groups demonstrated significantly higher SCR amplitude during the early compared to the late part of the phase. The levodopa group demonstrated significantly higher SCR compared to the placebo group, but no significant difference comparing stimulus types (CS+ or CS-; *Figure 2–B*). Non-parametric ATS revealed a significant main effect of Block (F_(1,65.5)_=34.23, p<0.001) and Group (F_(1.99,67)_=3.00, p<0.050), but no Stimulus difference (p=0.121). The Stimulus x Block (F_(1,67.3)_=7.6, p = 0.006) interaction was significant, all other interactions (all p>0.180) were not significant *(Table S6*). Post-hoc analysis of Stimulus x Block interactions demonstrated significantly higher CS+ compared to CS- SCR amplitude in early block only (p=0.016; late block

p=0.994) as well as higher CS+ and CS- SCR amplitude in early compared to late blocks (all p<0.008). Furthermore, overall, SCR amplitude was significantly higher during the task for the levodopa group compared to the placebo group (p=0.029), but not for the tiapride group compared to the placebo (p=0.275) or to the levodopa group (p=0.581). Reanalysis considering the first trial of recall only revealed no significant group differences (*Figure S2-B; Table S7*).

## Results group B

### Questionnaires

Across all group B participants, CS+ was rated significantly higher in unpleasantness, arousal, and fear compared to CS- following fear acquisition training, and these differences persisted through extinction and recall phases (*Figure S3; Table S8; Table S9*). Participants estimated a high likelihood of US occurrence following CS+ (96.40 ± 12.80%), while the probability was minimal for CS- (3.87% ± 15.24%; *Table S5)*.

### Pupillometry

#### Habituation phase

During the habituation phase, the mean (differential) pupil size towards CS+ and CS- did not differ between groups. Non-parametric ATS revealed no significant main effects of Stimulus (p=0.377), Group (p=0.643), or Stimulus x Group (p=0.626) interactions (*Figure 3-A; Table S10*).

**Figure 3.**
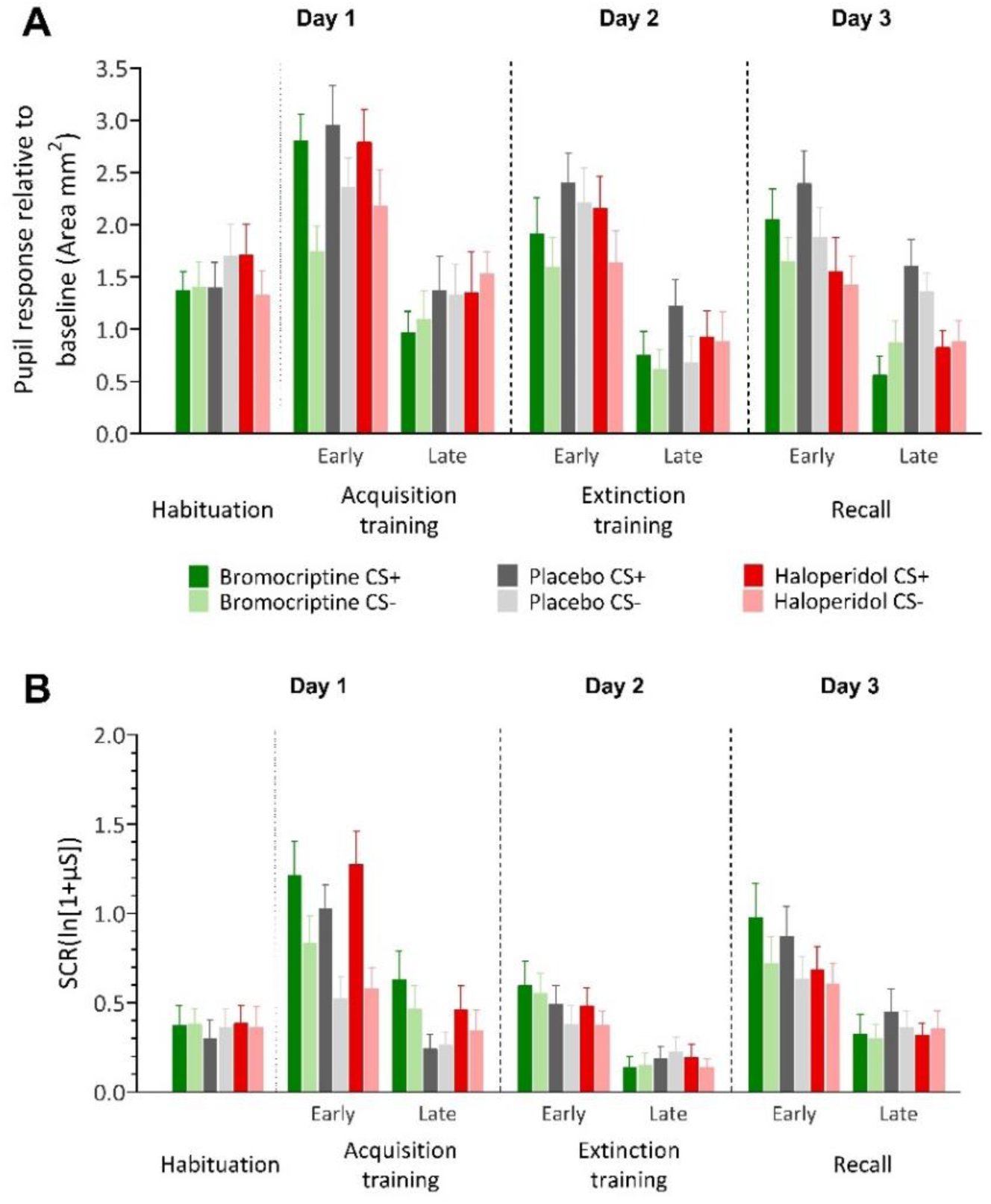
(A) **Pupil responses relative to baseline** for habituation and early and late blocks of fear acquisition and extinction training as well as the recall phase in the groups receiving bromocriptine, placebo or haloperidol. Bars represent means, error bars indicate S.E.M. (B) **Skin conductance responses** (SCRs) during habituation and early and late blocks of fear acquisition and extinction training as well as the recall phase in the groups receiving bromocriptine, placebo or haloperidol. Bars represent means, error bars indicate S.E.M.

#### Fear acquisition training

During fear acquisition training, bromocriptine, placebo and haloperidol groups showed significantly higher pupil responses to CS+ compared to CS- and during the early compared to the late part of the phase (*Figure 3-A*). Non-parametric ATS revealed a significant main effect of Stimulus (F_(1,66.6)_=6.84, p<0.009) and Block (F_(1,63.4)_=62.51, p<0.001), but no Group difference (p=0.602). Stimulus x Block (F_(1,69.4)_=15.36, p<0.001) interaction showed significant differences, other interactions (all p>0.105) were not significant (*Table S10*). Post-hoc analysis of Stimulus x Block demonstrated significantly higher CS+ and CS- pupil responses in early compared to late blocks (all p<0.001), and higher pupil responses for CS+ than for CS- trials in early (p<0.001) but not late blocks (p=0.978).

#### Extinction training

During extinction training, bromocriptine, placebo and haloperidol groups showed significantly higher pupil responses to CS+ compared to CS- and during the early compared to the late part of the phase (*Figure 3-A*). Non-parametric ATS revealed a significant main effect of Stimulus (F_(1,66.9)_=6.10, p<0.014) and Block (F_(1,68.5)_=69.26, p<0.001), but no Group difference (p=0.248). Additionally, none of the interactions were significant (Stimulus x Block (p=0.487), Group x Stimulus (p=0.560), Group × Block (p = 0.377) or Group x Stimulus x Block (p=0.278; *Table S10)*.

#### Recall

During recall, bromocriptine, placebo and haloperidol groups demonstrated significantly higher pupil size response during the early compared to the late part of the phase and significant differences between both groups and the placebo group, but no significant difference between stimulus types (CS+ or CS-) (*Figure 3-A).* Non-parametric ATS revealed a main effect of group (F_(1.98,69.8)_=4.98, p=0.007) and early compared to late blocks (F_(1,63.1)=_48.97, p<0.001), but no main effects for stimulus (p=0.716, *Table S10*). The Block x Group (F_(1.9,63.1)_=3.04, p=0.050) interaction was significant, all other interactions (all p>0.070) were not significant *(Table S10*). Post hoc analysis of the Block x Group interaction revealed significantly higher pupil responses in the early phase compared to the late phase for bromocriptine (p<0.001), placebo (p<0.009) and haloperidol groups (p<0.014). Pupil responses during the early block did not differ significantly between groups (all p>0.08). However, during the late block the pupil responses were lower in the bromocriptine compared to placebo group (p=0.009), and in the haloperidol compared to the placebo group (p=0.014). Reanalysis considering the first trial of recall only revealed no significant group differences (*Figure S4–A; Table S11*).

### Skin conductance responses (SCRs)

#### Habituation phase

During the habituation phase, the mean SCR amplitudes towards CS+ and CS- did not differ between groups (*Figure 3-B*). Non-parametric ATS revealed no significant main effects of Stimulus (p*=*0.387), Group (p=0.858), or Stimulus x Group (p=0.349) interactions (*Table S10)*.

#### Fear acquisition training

During fear acquisition training, bromocriptine, placebo and haloperidol groups showed significantly higher mean SCR amplitudes to CS+ compared to CS- and during the early compared to the late part of the phase (*Figure 3-B*). Non-parametric ATS revealed a significant main effect of Stimulus (F_(1,52.3)_=31.17, p<0.001) and Block (F_(1,70.1)_=70.96, p<0.001), but no Group effect (p=0.74). Both Stimulus x Block (F_(1,63)_=17.95, p<0.001) and Stimulus x Group x Block (F_(1.87,63)_=5.00, p<0.008) interactions showed significant differences, all other interactions (all p>0.41) were not significant (*Table S10*). Stimulus x Block x Group post hoc analysis revealed significantly higher SCR amplitudes during early CS+ compared to late CS+ for all three groups (all p<0.006), as well as in the early CS+ compared to early CS- (all p<0.016), but not during late part of the phase (all p>0.26). In both the bromocriptine and placebo groups, early CS- exhibited significantly higher SCR amplitudes than late CS- (all p<0.007). The haloperidol group did not show a significant decrease in SCR amplitude levels between early and late CS- (p=0.503). Compared to the placebo group, no significant differences were observed for haloperidol (all p>0.67) and bromocriptine (all p>0.867) regarding both CS+ and CS- SCR amplitudes during either early or late parts of the phase.

#### Extinction training

During extinction training, all three groups demonstrated significantly higher SCR amplitude during the early compared to the late part of the phase with no significant differences between groups and group stimulus types (CS+ vs. CS-) (*Figure 3-B*). Non-parametric ATS revealed a significant main effect of block (F_(1,71.4)_=32.73, p<0.001), while no main effects were observed for Stimulus (p=0.280) and Group (p=0.916). Additionally, none of the interactions (Stimulus x Block (p=0.302), Group x Stimulus (p=0.860), Group × Block (p=0.218), and Group x Stimulus x Block (p=0.699)) were significant (*Table S10)*.

#### Recall

During recall, all three groups demonstrated significantly higher SCR amplitude during the early compared to the late part of the phase. There was no significant difference comparing groups or stimulus types (CS+ or CS-; *Figure 3-B*). Non-parametric ATS revealed a significant main effect of block (F_(1,61.4)_=57.08, p<0.001), while no main effects were observed for Stimulus (p=0.100) and Group (p=0.960). Additionally, none of the interactions were significant (Stimulus x Block (p= 0.114), Group x Stimulus (p= 0.682), Group × Block (p= 0.291) or Group x Stimulus x Block (p= 0.594; *Table S10)*. Reanalysis considering the first trial of recall only revealed no significant group differences (*Figure S4–B; Table S11*).

## Baseline

In the baseline pupil size analysis (prestimulus, that is, 2s prior to CS onset; *Figure 4*), the placebo groups (A and B) showed no significant differences between phases (p=0.357 for Group A, p=0.571 for Group B). Dopamine agonists, levodopa and bromocriptine, showed significant differences between only two phases: levodopa had a higher baseline during fear acquisition compared to extinction (F_(2.99,728)_=4.38, p=0.004), and bromocriptine had a higher baseline during extinction compared to recall (F_(3,754)_=3.03, p=0.028) (*Table S12*). These limited differences suggest that the effects in the dopamine agonist groups may not be directly attributed to the drug. In contrast, dopamine antagonists tiapride and haloperidol showed significant decreases in baseline pupil size during one phase compared to all others: tiapride during extinction (F_(3,682)_=22.94, p<0.001) and haloperidol during recall (F_(3,794)_=5.64, p<0.001). These reductions in pupil size corresponded with the presence of the drugs in the blood, indicating intrinsic effects of the dopamine antagonists. For tiapride, which has a short half-life, the reduction in pupil size is observed during extinction training (*Table S2*). For haloperidol, the longer half-life of haloperidol, evidenced by the higher drug concentrations in the blood during recall compared to extinction, likely explains this observation (*Table S2).* Note that no bromocriptine level was measured on day 2, likely due to the sensitivity limitations of the available test (detection limit: 0.1 ng/mL).

**Figure 4.**
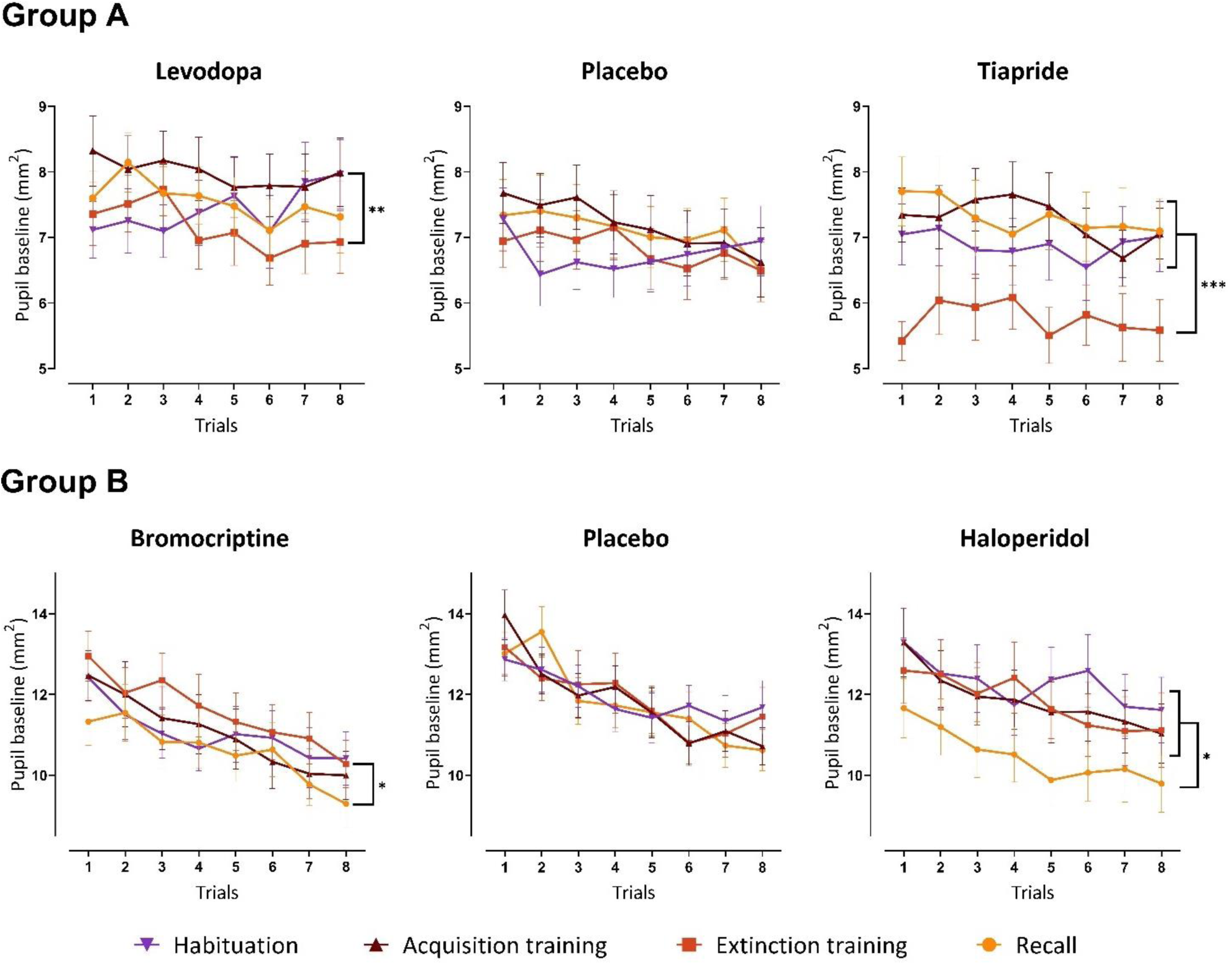
Pupil size variation (in mm^2^) 2s prior to CS onset during the first 8 trials in both group A receiving levodopa, placebo or tiapride and group B receiving bromocriptine, placebo or haloperidol. All data shown as means and S.E.M. (*p<0.05; **p<0.01; *** p<0.001)

## Discussion

The main aim of the study was to explore the effect of dopaminergic agonists and antagonists on extinction learning and subsequent extinction and recall in humans, as measured by changes in autonomic conditioned responses (SCR and pupil size) and self-reported questionnaires outcomes. Findings provide some additional support that the dopamine system is involved in fear extinction learning. Drug effects, however, were small and unspecific drug effects were also observed.

### Dopaminergic effects on fear extinction learning

We tested the hypothesis that pharmacological intake of different dopaminergic drugs (levodopa and bromocriptine) prior to extinction would enhance fear extinction learning, leading to a reduction in fear recall the following day. We were thus expecting to observe a diminution of SCR and pupil size conditioned responses during both extinction training and the recall phase for both levodopa and bromocriptine groups compared to their respective placebo. However, although we found effects of dopaminergic drugs, findings were present during recall only, and effects following bromocriptine and levodopa intake were opposing.

In line with our hypothesis, we measured reduced pupil dilation during late recall in the bromocriptine group compared to the placebo group, indicating faster extinction of spontaneously recovered fear reactions on the third day. This could suggest that bromocriptine, as a dopaminergic agonist acting on postsynaptic dopamine D2 receptors, may facilitate fear extinction consolidation, thereby allowing for faster re-extinction on the following day. This aligns with previous findings where single doses of bromocriptine (1.25 or 2.5 mg) have demonstrated improvement in learning and memory formation in reversal learning, working memory and cognitive flexibility^21,33,46,47^.

On the other hand, contrary to our expectations, the levodopa group showed significantly higher SCR amplitudes during spontaneous recovery compared to the placebo group. This unexpected result may be explained by the intrinsic individual differences in SCR arousal levels^42^, as supported by the fact that during habituation phase, before any drug intake, participants in levodopa group already demonstrated higher overall SCR amplitudes. Furthermore, findings from Andres et al.,^48^ suggest that the effect of levodopa in enhancing extinction memory consolidation may be favoured by environmental stress/arousal induce by an MRI. Indeed, they reported pronounced levodopa effects during experiments conducted within MRI environments^19,22^. However, when experiments were behavioral only, conditional effects of levodopa were observed on SCR measures only in selected participants who had demonstrated successful extinction learning^23^.

### Anti-dopaminergic effects on fear extinction learning

We also investigated whether administering distinct anti-dopaminergic drugs (tiapride and haloperidol) prior to extinction would hinder fear response modulation during both fear extinction training and subsequent recall. Following the same logic, we anticipated less reduction of SCR and pupil size conditioned responses during both extinction training and recall for both tiapride and haloperidol groups. However, unexpectedly, tiapride intake did not seem to affect fear extinction and haloperidol demonstrated a decrease in pupil CR in late recall.

Additionally, contrary to our expectations, the group who took haloperidol showed no difference in SCRs and a diminished pupillary reaction during the late recall phase compared to its placebo group, suggesting a potential reduction in conditioned responses (CR). This is surprising given previous findings by Holtzman-Assif et al.^18^ showing that haloperidol-injected rats displayed increasing freezing behavior compared to a control group during extinction sessions and on a drug-free recall test, indicating an impairment in fear response inhibition. Moreover, this effect was particularly pronounced when haloperidol was infused in the nucleus accumbens, underscoring its critical role in fear extinction learning.

Interestingly, both dopamine antagonists appeared to have an intrinsic effect on the baseline tonic pupil size measure prior each stimulus. We found that tiapride intake resulted in a reduction of baseline pupil size levels during extinction training compared to habituation, fear acquisition training, and recall. Similarly, haloperidol intake led to a reduction of baseline levels during the recall phase compared to habituation, fear acquisition training, and extinction training. Given that the drug is administered before extinction training, the reduction in baseline observed during recall but not extinction training following haloperidol intake may initially seem counterintuitive. However, this could be attributed to the prolonged half-life of haloperidol. The analysis of blood concentrations confirmed a higher presence of haloperidol in the blood on day 3 compared to day 2, all the other drugs were only detected on day 2. According to the literature, the tonic pupil size impacts the magnitude of the following pupil responses^26^. The effect of tiapride and haloperidol intake on tonic pupil size might explain the unexpected decrease in pupillary conditioned responses, contrary to the anticipated increase.

### Limitations

The present study shows that assessing the dopaminergic system in healthy human participants by administration of oral dopaminergic and anti-dopaminergic drugs has limitations. One of the challenges in understanding the involvement of dopamine in fear extinction learning in humans is based on the widespread distribution of dopamine receptors across multiple areas of the brain. Each region, from the VTA to the nucleus accumbens, including amygdala, hippocampus, and prefrontal cortex, is thought to play a part in fear extinction, with dopamine likely exerting differential effects depending on the receptor subtype, dose and specific neural circuit involved.

By orally administering drugs that modulate the overall dopamine levels, contradictory effects may be observed in brain regions containing dopamine receptors^49^, leading to a result which is not clearly enhancing or impairing fear extinction learning.

Although the chosen timing and dosage of the respective drugs were based on existing literature, it did not consider individual variability. While it may be difficult and costly to try to evaluate individual’s pharmacokinetics prior to the experiment, one could consider personalized dosing based on sex, age and weight. Furthermore, drug dosages may need to be higher to achieve significant effects. For example, based on the analytical technique used, we did not observe bromocriptine in blood samples taken after the extinction phase on day 2.

Consistent with the existing literature, our study observed clear differential conditioned responses for SCRs^26,28,29^ and pupil size^26,28^. However, while SCRs, pupil responses and self-reported fear questionnaires can provide an overall assessment of fear acquisition and fear extinction success, it may not have been sufficient to measure the subtle change in conditioned responses attributable to the effects of dopaminergic modulation. Future studies could benefit from brain imaging studies, in particular combining PET/MRI scanner^50^ to monitor dopamine release during extinction training and evaluate the changes due to systematic dopamine agonist or antagonist drug intake.

Finally, group sizes may have been too small. We observed differences between the groups already during habituation and fear acquisition training, that is prior drug intake, which is likely explained by individual variability in physiological measures.

### Summary and conclusions

In sum, bromocriptine intake resulted in faster re-extinction during recall, suggesting more robust consolidation of fear extinction. Contrary to our expectations, levodopa resulted in increased spontaneous recovery during early recall, possibly due to individual differences in SCR arousal levels. Regarding dopamine antagonists, tiapride showed no significant effects on fear extinction learning, while haloperidol unexpectedly led to faster re-extinction during late recall, potentially due to its prolonged half-life and intrinsic effects on tonic pupil size. Overall while dopaminergic drugs show potential in modulating fear extinction and recall, providing support to the fact that the dopaminergic system contributes to extinction learning in humans, their effects are complex to interpret and seem to be influenced by multiple factors, including individual variability, drug receptor and environmental context.

## Data availability

The data that support the findings of this study are available on request from the corresponding author.

## Acknowledgements

We thank Greta Wippich for making the experimental set-up illustration and Beate Brol for supporting with participants coordination.

## Author contributions

Data acquisition was performed by A.D., L.M. and K.K., supported by A.T., B.A., S.N., F.E. and D.T. Data preprocessing was performed by A.D., L.M., K.K. and E.N.. Data analysis was performed by A.D., G.B. and T.E.. D.T., C.J.M. S.C., T.E. designed and supervised the study. A.D. wrote the manuscript, all authors read, edited and approved the manuscript. All authors had complete access to the study data that support the publication.

---

**Table S1.**
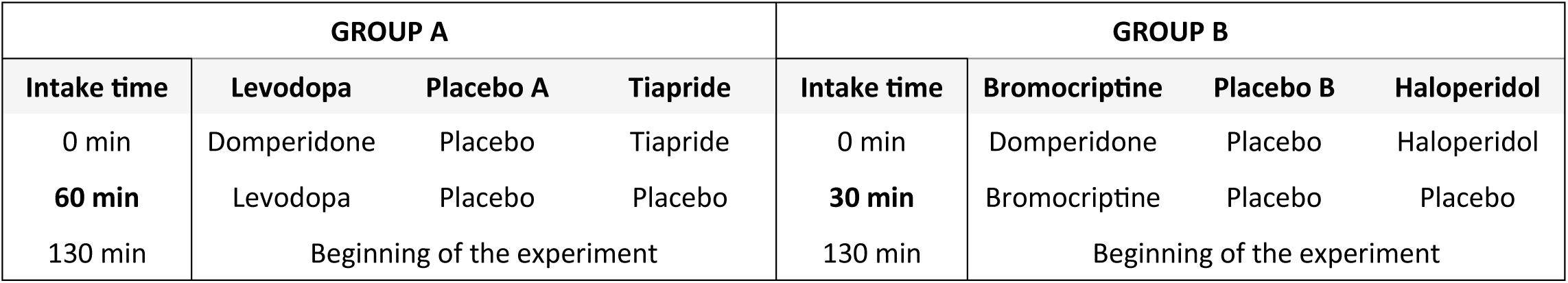
Medication protocols of the different dopaminergic and anti-dopaminergic drugs day 2.

| GROUP A |  |  |  | GROUP B |  |  |  |
| --- | --- | --- | --- | --- | --- | --- | --- |
| Intake time | Levodopa | Placebo A | Tiapride | Intake time | Bromocriptine | Placebo B | Haloperidol |
| 0 min | Domperidone | Placebo | Tiapride | 0 min | Domperidone | Placebo | Haloperidol |
| 60 min | Levodopa | Placebo | Placebo | 30 min | Bromocriptine | Placebo | Placebo |
| 130 min | Beginning of the experiment |  |  | 130 min | Beginning of the experiment |  |  |

**Table S2.** Drug concentration measured each day in the participants blood after the experiment. * below detection limit 0.1 ng/mL. Measurements conducted by **MVZ Medizinisches Labor Bremen GmbH and by ***MVZ Dr. Eberhard & Partner Dortmund.

| Drugs | Blood level |  |  |
| --- | --- | --- | --- |
|  | Day 1 | Day 2 | Day 3 |
| Placebo A (n=25) | - | - | - |
| Levodopa (100 mg) (n=25) *** | - | 0.66 ± 0.34 (µg/mL) | - |
| Tiapride (100 mg) (n=22) *** | - | 0.54 ± 0.11 (µg/mL) | - |
| Placebo B (n=25) | - | - | - |
| Bromocriptine (1.25 mg) (n=25) ** | - | * | - |
| Haloperidol (3 mg) (n=22) *** | - | 0.24 ± 0.43 (ng/mL) | 0.35 ± 0.22 (ng/mL) |

## Supplementary materials – Group A

### Questionnaires

**Figure S1.**
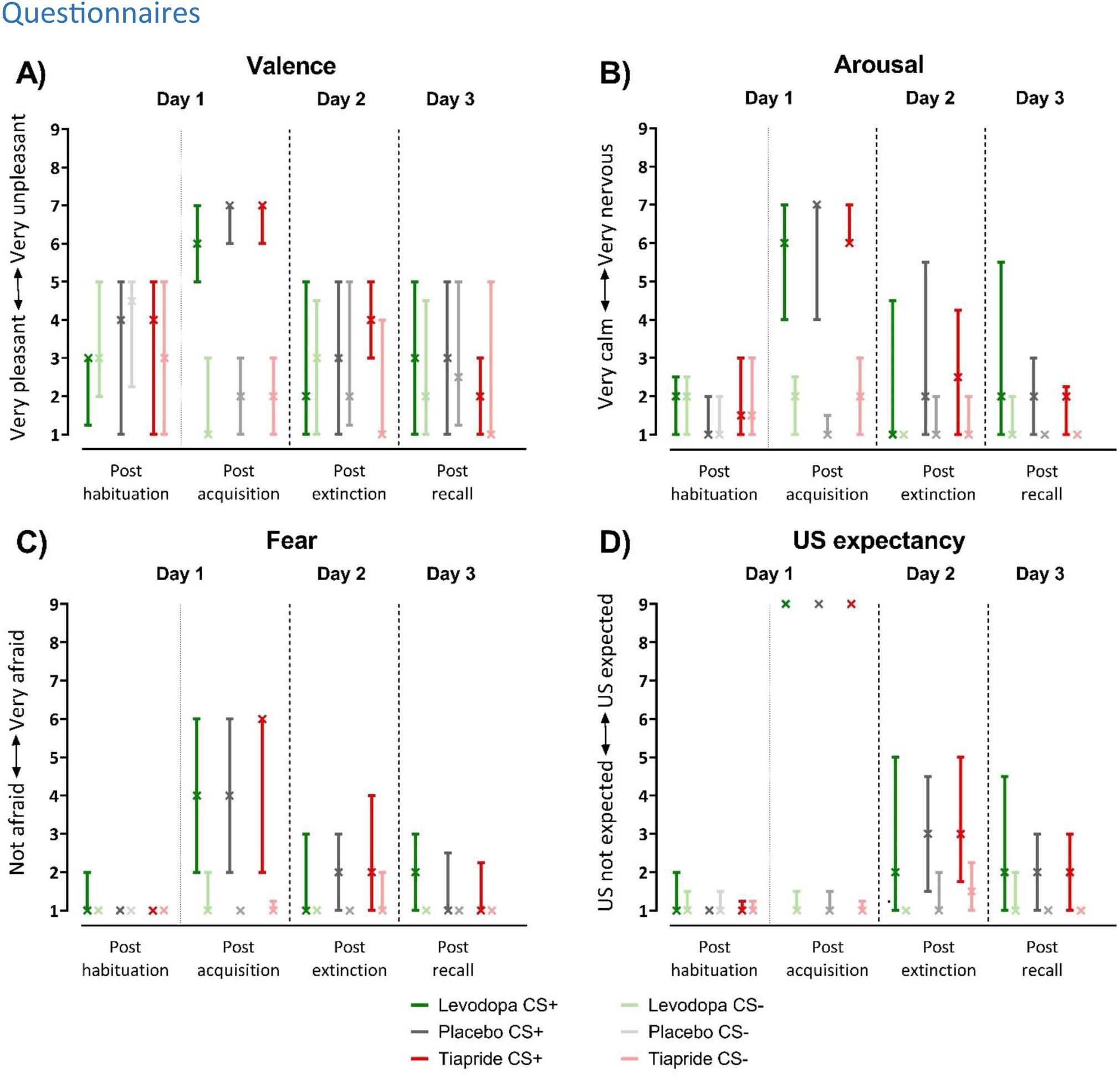
Self-reported fear evaluation questionnaire ratings in the groups receiving levodopa, placebo or tiapride post habituation, fear acquisition training, extinction training, and recall. The median is marked by a cross (whiskers extend to the 1st and 3rd quartiles) for ratings of valence (A), arousal (B), fear (C), and US expectancy (D) based on a 1-9 Likert scale.

#### CS valence, arousal, and fear ratings

Following the habituation phase no significant differences in valence, arousal, and fear ratings between CS+ and CS- were present, and this was consistent across levodopa, placebo and tiapride groups. After fear acquisition training, CS+ received significantly higher unpleasantness ratings, higher arousal ratings, and was perceived as inducing more fear than CS- in all three groups (*Figure S1*). These distinctions between CSs persisted in every group during extinction and recall phases. Post acquisition phase ratings consistently highlighted significant differences between CS+ and CS- in valence, arousal and fear ratings in all groups (*Table S3)*. Non-parametric ANOVA-type statistic revealed a significant main effect of Stimulus and Phase (main effects: all p<0.001), and a Stimulus × Phase interaction for arousal (F_(2.71,164)=_40.30, p<0.001), valence (F_(2.51,76.2)=_37.91, p<0.001) and fear (F_(2.69,153)=_34.91, p<0.001) ratings independently of the drug group (*Table S4*). Post-hoc tests showed significant differences between stimuli post fear acquisition training, extinction training and recall phases (least square means tests, Valence: all p<0.001; Arousal: all p<0.001; Fear: all p<0.001), but not following the habituation phase (least square means test, all p>0.80). No significant differences in arousal, fear and valence ratings between groups were found.

#### US unpleasantness, CS/US contingency and US expectancy

Across all participants of the three groups, the likelihood that a US onset occurred after CS+ presentation was estimated to be 98.21 ± 11.40% (100% probability for 68 out of 71 participants), while after CS- presentation it was 1.54 ± 5.34% (0% probability for 69 out of 71 participants; *Table S5*). Overall median US unpleasantness was rated 7 (IQR 7-8) following acquisition, participants reported recognizing the association between CS+ and US after experiencing an average of 2.52 ± 1.63 electric shocks. No significant between groups for both US unpleasantness and CS/US contingency ratings have been found.

After habituation phase, reported US expectancy following CS+ and CS- were not significantly different from each other. Post fear acquisition training, participants reported a higher US expectancy after CS + compared to the CS- and this difference remained until the end of recall. Similarly to valence, arousal and fear ratings, post-acquisition phase consistently showed significant differences between CS+ and CS- in US expectancy ratings in all groups (*Figure S1; Table S3*). Non-parametric ANOVA-type statistic revealed a significant main effect of Stimulus and Phase (main effects: all p<0.001), and a Stimulus × Phase interaction for US expectancy (F_(2.39,155)_=136.24, p<0.001) ratings independently of the drug group (*Table S4*). Post-hoc tests showed significant differences between stimuli post fear acquisition training, extinction training and recall (least square means tests, US expectancy: all p<0.001), but not following habituation (least square means test, p>0.96).

**Table S3.**
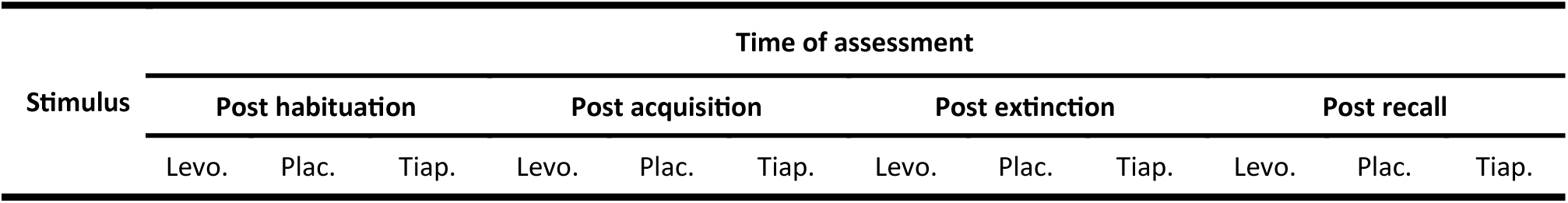
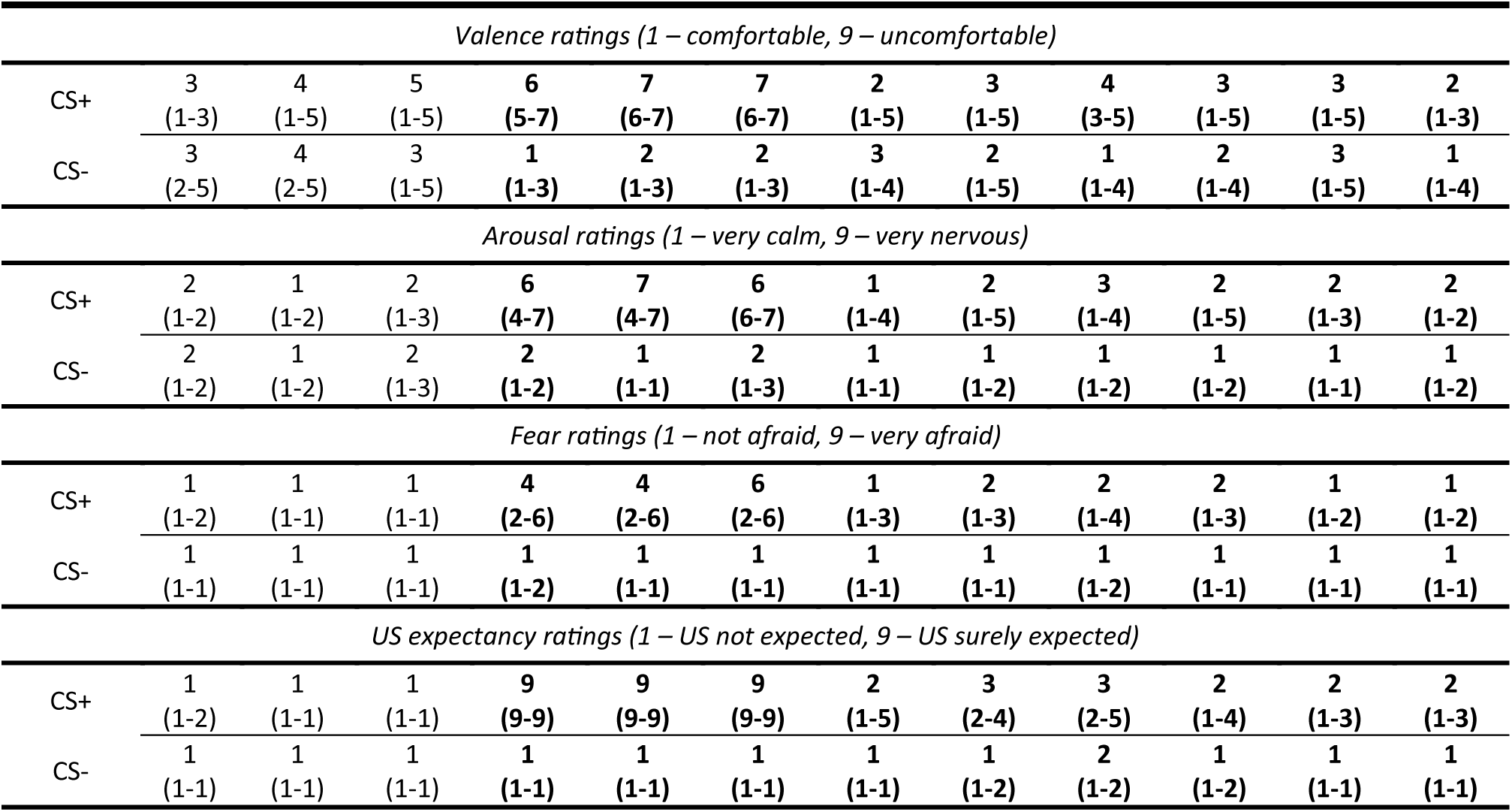
Self-reported fear evaluation questionnaire ratings in the groups receiving levodopa (Levo.), placebo (Plac.) or tiapride (Tiap.) post habituation, fear acquisition training, extinction training, and recall. Median (interquartile range) ratings of valence (A), arousal (B), fear (C) and US expectancy (D) based on a 1-9 Likert scale. Statistically significant differences between CS+ and CS- are shown in bold (least square means tests, p<0.01). Note that no statistically significant differences were found between drug groups and placebo group.

| Stimulus | Time of assessment |  |  |  |  |  |  |  |  |  |  |  |
| --- | --- | --- | --- | --- | --- | --- | --- | --- | --- | --- | --- | --- |
|  | Post habituation |  |  | Post acquisition |  |  | Post extinction |  |  | Post recall |  |  |
|  | Levo. | Plac. | Tiap. | Levo. | Plac. | Tiap. | Levo. | Plac. | Tiap. | Levo. | Plac. | Tiap. |

| *Valence ratings (1 – comfortable, 9 – uncomfortable)* |  |  |  |  |  |  |  |  |  |  |  |  |
| --- | --- | --- | --- | --- | --- | --- | --- | --- | --- | --- | --- | --- |
| CS+ | 3<br>(1-3) | 4<br>(1-5) | 5<br>(1-5) | 6<br>(5-7) | 7<br>(6-7) | 7<br>(6-7) | 2<br>(1-5) | 3<br>(1-5) | 4<br>(3-5) | 3<br>(1-5) | 3<br>(1-5) | 2<br>(1-3) |
| CS- | 3<br>(2-5) | 4<br>(2-5) | 3<br>(1-5) | 1<br>(1-3) | 2<br>(1-3) | 2<br>(1-3) | 3<br>(1-4) | 2<br>(1-5) | 1<br>(1-4) | 2<br>(1-4) | 3<br>(1-5) | 1<br>(1-4) |
| *Arousal ratings (1 – very calm, 9 – very nervous)* |  |  |  |  |  |  |  |  |  |  |  |  |
| CS+ | 2<br>(1-2) | 1<br>(1-2) | 2<br>(1-3) | 6<br>(4-7) | 7<br>(4-7) | 6<br>(6-7) | 1<br>(1-4) | 2<br>(1-5) | 3<br>(1-4) | 2<br>(1-5) | 2<br>(1-3) | 2<br>(1-2) |
| CS- | 2<br>(1-2) | 1<br>(1-2) | 2<br>(1-3) | 2<br>(1-2) | 1<br>(1-1) | 2<br>(1-3) | 1<br>(1-1) | 1<br>(1-2) | 1<br>(1-2) | 1<br>(1-2) | 1<br>(1-1) | 1<br>(1-2) |
| *Fear ratings (1 – not afraid, 9 – very afraid)* |  |  |  |  |  |  |  |  |  |  |  |  |
| CS+ | 1<br>(1-2) | 1<br>(1-1) | 1<br>(1-1) | 4<br>(2-6) | 4<br>(2-6) | 6<br>(2-6) | 1<br>(1-3) | 2<br>(1-3) | 2<br>(1-4) | 2<br>(1-3) | 1<br>(1-2) | 1<br>(1-2) |
| CS- | 1<br>(1-1) | 1<br>(1-1) | 1<br>(1-1) | 1<br>(1-2) | 1<br>(1-1) | 1<br>(1-1) | 1<br>(1-1) | 1<br>(1-1) | 1<br>(1-2) | 1<br>(1-1) | 1<br>(1-1) | 1<br>(1-1) |
| *US expectancy ratings (1 – US not expected, 9 – US surely expected)* |  |  |  |  |  |  |  |  |  |  |  |  |
| CS+ | 1<br>(1-2) | 1<br>(1-1) | 1<br>(1-1) | 9<br>(9-9) | 9<br>(9-9) | 9<br>(9-9) | 2<br>(1-5) | 3<br>(2-4) | 3<br>(2-5) | 2<br>(1-4) | 2<br>(1-3) | 2<br>(1-3) |
| CS- | 1<br>(1-1) | 1<br>(1-1) | 1<br>(1-1) | 1<br>(1-1) | 1<br>(1-1) | 1<br>(1-1) | 1<br>(1-1) | 1<br>(1-2) | 2<br>(1-2) | 1<br>(1-2) | 1<br>(1-1) | 1<br>(1-1) |

**Table S4.** Self-reported fear evaluation questionnaire ratings in the groups receiving levodopa, placebo or tiapride. Results of the non-parametric ANOVA-type statistics for repeated measures on all phases.

| Factor | Num DF | Den DF | F | Pr > F(DDF) | Pr>F(infty) |
| --- | --- | --- | --- | --- | --- |
| Valence |  |  |  |  |  |
| Phase | 2.49 | 72.2 | 16.38 | <.001 *** | <.001 *** |
| Stimulus | 1 | 57.3 | 29.59 | <.001 *** | <.001 *** |
| Group | 1.98 | 67.5 | 0.41 | 0.668 | 0.663 |
| Stimulus * Phase | 2.51 | 76.2 | 37.91 | <.001 *** | <.001 *** |
| Group * Phase | 4.6 | 72.2 | 0.78 | 0.558 | 0.554 |
| Stimulus * Group | 1.99 | 57.3 | 0.84 | 0.435 | 0.430 |
| Stimulus * Group * Phase | 4.66 | 76.2 | 0.37 | 0.855 | 0.856 |
| Arousal |  |  |  |  |  |
| Phase | 2.5 | 154 | 26.80 | <.001 *** | <.001 *** |
| Stimulus | 1 | 67.9 | 162.16 | <.001 *** | <.001 *** |
| Group | 1.99 | 67 | 0.54 | 0.587 | 0.584 |
| Stimulus * Phase | 2.71 | 164 | 40.30 | <.001 *** | <.001 *** |
| Group * Phase | 4.78 | 154 | 1.62 | 0.162 | 0.155 |
| Stimulus * Group | 2 | 67.9 | 1.15 | 0.324 | 0.318 |
| Stimulus * Group * Phase | 5.16 | 164 | 0.34 | 0.895 | 0.896 |
| Fear |  |  |  |  |  |
| Phase | 2.74 | 171 | 29.76 | <.001 *** | <.001 *** |
| Stimulus | 1 | 67.4 | 146.39 | <.001 *** | <.001 *** |
| Group | 2 | 67.9 | 0.13 | 0.881 | 0.881 |
| Stimulus * Phase | 2.69 | 153 | 34.91 | <.001 *** | <.001 *** |
| Group * Phase | 5.25 | 171 | 1.50 | 0.190 | 0.183 |
| Stimulus * Group | 1.99 | 67.4 | 0.14 | 0.865 | 0.865 |
| Stimulus * Group * Phase | 4.97 | 153 | 0.50 | 0.772 | 0.773 |
| US Expectancy |  |  |  |  |  |
| Phase | 2.7 | 204 | 39.29 | <.001 *** | <.001 *** |
| Stimulus | 1 | 64.9 | 374.69 | <.001 *** | <.001 *** |
| Group | 1.91 | 101 | 0.30 | 0.728 | 0.727 |
| Stimulus * Phase | 2.39 | 155 | 136.24 | <.001 *** | <.001 *** |
| Group * Phase | 5.15 | 204 | 1.10 | 0.360 | 0.356 |
| Stimulus * Group | 1.96 | 64.9 | 0.18 | 0.832 | 0.831 |
| Stimulus * Group * Phase | 4.64 | 155 | 0.64 | 0.661 | 0.660 |

**Table S5.** Fear conditioning questionnaires on US in both group A receiving levodopa, placebo or tiapride and group B receiving bromocriptine, placebo or haloperidol. Median (interquartile range) of US expectancy ratings and mean percentage of US perception following each CS and number of US perceived before CS/US contingency recognition post fear acquisition. Note that no statistically significant differences were found between drug groups and placebo group.

| Post acquisition |  |  |  |  |  |  |
| --- | --- | --- | --- | --- | --- | --- |
|  | Levodopa | Placebo | Tiapride | Bromocriptine | Placebo | Haloperidol |
| *US unpleasantness (1 – US not unpleasant, 9 – US highly unpleasant)* |  |  |  |  |  |  |
| US | 7 (6-8) | 7 (7-8) | 8 (7-8) | 8 (7-8) | 7 (6-8) | 8 (7-8) |
| Estimated probability of CS followed by US (%) |  |  |  |  |  |  |
| CS+ | 100.00<br>100 % for 24/24 | 96.80 ± 16.00<br>100 % for 24/25 | 97.73 ± 8.69<br>100 % for 20/22 | 97.20 ± 14.00<br>100 % for 24/25 | 94.80 ± 14.18<br>100 % for 21/25 | 97.20 ± 10.21<br>100 % for 22/25 |
| CS- | 0.00<br>0% for 24/24 | 0.40 ± 2.00<br>0% for 24/25 | 2.27 ± 8.69<br>0% for 20/22 | 1.20 ± 6.00<br>0% for 24/25 | 8.00 ± 22.55<br>0% for 22/25 | 2.40 ± 12.00<br>0% for 24/25 |
| *Number of US perceived before CS/US contingency recognition* |  |  |  |  |  |  |
| CS/US | 2.80 ± 1.38 | 2.2 ± 0.96 | 2.59 ± 1.84 | 2.13 ± 0.80 | 2.38 ± 1.53 | 2.30 ± 1.82 |

### Statistics for pupillometry and skin conductance responses

**Table S6.** Pupil size and skin conductance responses statistics for the groups receiving levodopa, placebo or tiapride. Results of the non -parametric ANOVA-type statistics for repeated measures on all phases. (*p<0.01; ** p<0.05; *** p<0.001)

| Pupillometry |  |  |  |  |
| --- | --- | --- | --- | --- |
| Factor | Num DF | Den DF | F | Pr>F(nifty ) |
| Habituation |  |  |  |  |
| Stimulus | 1 | 60.4 | 1.42 | 0.233 |
| Group | 1.95 | 61.4 | 0.95 | 0.383 |
| Stimulus * Group | 1.84 | 60.4 | 0.46 | 0.613 |
| Fear Acquisition training |  |  |  |  |
| Block | 1 | 66.7 | 32.25 | **<.001 \*\*\*** |
| Stimulus | 1 | 55.5 | 37.40 | **<.001 \*\*\*** |
| Group | 1.96 | 61.5 | 1.22 | 0.296 |
| Stimulus * Block | 1 | 65.1 | 1.48 | 0.224 |
| Group * Block | 1.95 | 66.7 | 2.96 | 0.053 |
| Stimulus * Group | 1.86 | 55.5 | 3.22 | **0.044 \*** |
| Stimulus * Group * Block | 1.96 | 65.1 | 3.30 | **0.038 \*** |
| Extinction training |  |  |  |  |
| Block | 1 | 61.3 | 33.80 | **<.001 \*\*\*** |
| Stimulus | 1 | 62.2 | 0.82 | 0.365 |
| Group | 2 | 64.5 | 2.82 | 0.060 |
| Stimulus * Block | 1 | 63.1 | 5.50 | **0.019 \*** |
| Group * Block | 1.96 | 61.3 | 0.95 | 0.384 |
| Stimulus * Group | 1.89 | 62.2 | 1.05 | 0.346 |
| Stimulus * Group * Block | 1.99 | 63.1 | 0.17 | 0.843 |
| Recall |  |  |  |  |
| Block | 1 | 52.2 | 15.19 | **<.001 \*\*\*** |
| Stimulus | 1 | 62.1 | 0.95 | 0.330 |
| Group | 1.99 | 63.5 | 2.04 | 0.130 |
| Stimulus * Block | 1 | 55.6 | 0.00 | 0.950 |
| Group * Block | 1.84 | 52.2 | 0.73 | 0.473 |
| Stimulus * Group | 1.99 | 62.1 | 0.13 | 0.879 |
| Stimulus * Group * Block | 1.88 | 55.6 | 0.56 | 0.562 |

| SCR |  |  |  |  |
| --- | --- | --- | --- | --- |
| Factor | Num DF | Den DF | F | Pr>F(nifty) |
| Habituation |  |  |  |  |
| Stimulus | 1 | 59.8 | 0.48 | 0.490 |
| Group | 1.99 | 67.1 | 2.58 | 0.076 |
| Stimulus * Group | 1.91 | 59.8 | 6.46 | **0.002 \*\*** |
| Fear Acquisition training |  |  |  |  |
| Block | 1 | 62.1 | 59.45 | **<.001 \*\*\*** |
| Stimulus | 1 | 67 | 65.10 | **<.001 \*\*\*** |
| Group | 1.97 | 64.4 | 2.74 | 0.066 |
| Stimulus * Block | 1 | 65.1 | 6.17 | **0.013 \*** |
| Group * Block | 1.93 | 62.1 | 0.14 | 0.859 |
| Stimulus * Group | 1.98 | 67 | 0.18 | 0.833 |
| Stimulus * Group * Block | 1.97 | 65.1 | 0.17 | 0.844 |
| Extinction training |  |  |  |  |
| Block | 1 | 60 | 65.60 | **<.001 \*\*\*** |
| Stimulus | 1 | 58.6 | 11.08 | **<.001 \*\*\*** |
| Group | 1.94 | 63.2 | 1.85 | 0.158 |
| Stimulus * Block | 1 | 63.1 | 4.75 | **0.029 \*** |
| Group * Block | 1.89 | 60 | 0.54 | 0.573 |
| Stimulus * Group | 1.86 | 58.6 | 0.28 | 0.741 |
| Stimulus * Group * Block | 1.9 | 63.1 | 0.57 | 0.554 |
| Recall |  |  |  |  |
| Block | 1 | 65.5 | 34.23 | **<.001 \*\*\*** |
| Stimulus | 1 | 50.4 | 2.40 | 0.121 |
| Group | 1.99 | 67 | 3.00 | **0.050 \*** |
| Stimulus * Block | 1 | 67.3 | 7.60 | **0.006 \*\*** |
| Group * Block | 1.96 | 65.5 | 1.06 | 0.345 |
| Stimulus * Group | 1.75 | 50.4 | 1.74 | 0.180 |
| Stimulus * Group * Block | 1.99 | 67.3 | 0.63 | 0.529 |

#### First recall trials pupillometry and skin conductance responses

**Figure S2.**
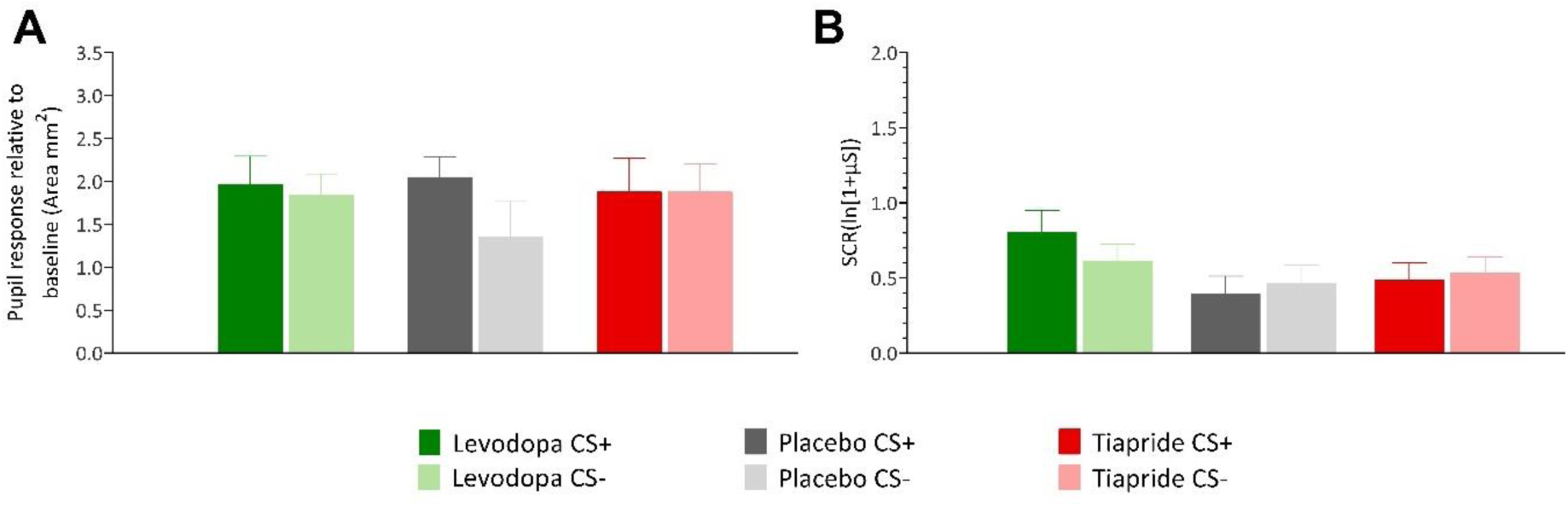
(A) ***Pupil responses relative to baseline*** *for first recall trial in the groups receiving levodopa, placebo or tiapride.* Bars represent means, e*rror bars indicate S.E.M. (B) **Skin conductance responses** (SCRs) for first recall trial in the groups receiving levodopa, placebo or tiapride.* Bars represent means, e*rror bars indicate S.E.M*.

We assessed the first trial in recall across levodopa, placebo, and tiapride groups using both pupil dilation and SCR (*Figure S2 – A and B; Table S7*). Both pupil dilation and SCR showed a significant difference between CS+ and CS- trials (Pupil: F(1, 55)=4.19, p=0.045; SCR: p<0.006), indicating spontaneous recovery of the initial fear association. However, there were no significant differences between the drug groups for either measure (Pupil: F(1.97, 59.5)=0.79, p=0.451; SCR: F(1.96, 63.6)=2.07, p=0.127). For the interaction between CS type and drug group, neither measure showed a significant effect (Pupil: F(1.95, 55)=1.47, p=0.230; SCR: F(1.91, 60.1)=2.55, p=0.081; *Table S7*).

**Table S7.** Pupillometry and SCR statistics in the groups receiving levodopa, placebo and tiapride. Results of the non-parametric ATS for repeated measures on the first trial of recall. (*p < 0.01; ** p < 0.05; *** p < 0.001)

| Factor | Num DF | Den DF | F | Pr > F(DDF) | Pr>F(infty) |
| --- | --- | --- | --- | --- | --- |
| Pupillometry |  |  |  |  |  |
| Stimulus | 1 | 55 | 4.19 | 0.045 | **0.041 \*** |
| Group | 1.97 | 59.5 | 0.79 | 0.456 | 0.451 |
| Stimulus * Group | 1.95 | 55 | 1.47 | 0.239 | 0.230 |
| SCR |  |  |  |  |  |
| Stimulus | 1 | 60.1 | 8.07 | 0.006 | **0.005 \*\*** |
| Group | 1.96 | 63.6 | 2.07 | 0.136 | 0.127 |
| Stimulus * Group | 1.91 | 60.1 | 2.55 | 0.089 | 0.081 |

## Supplementary materials – Group B

### Questionnaires

**Figure S3.**
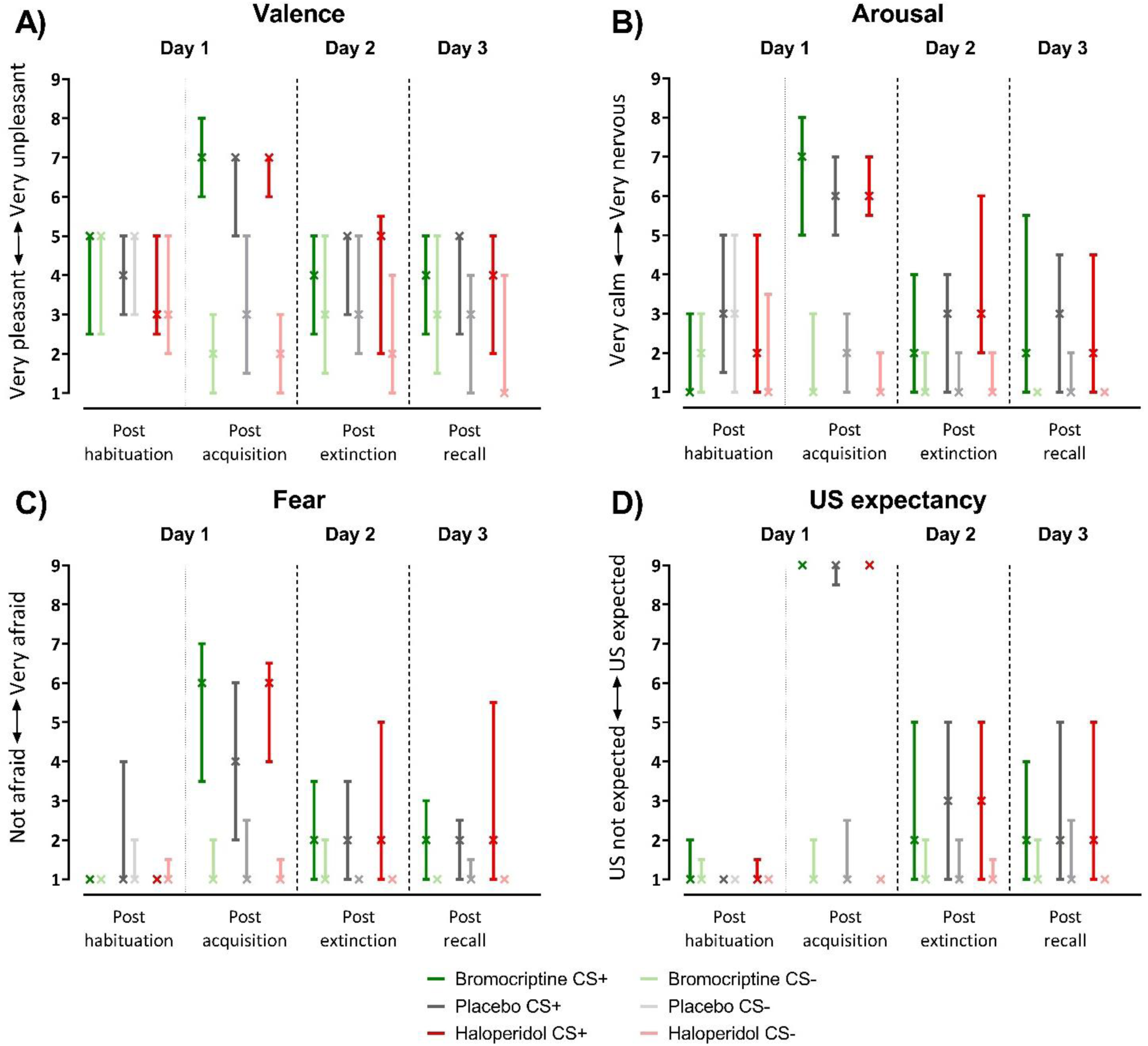
Self-reported fear evaluation questionnaire ratings in the groups receiving bromocriptine, placebo or haloperidol post habituation, fear acquisition training, extinction training, and recall. The median is marked by a cross (whiskers correspond to the 1st and 3rd quartiles) for ratings of valence (A), arousal (B), fear (C), and US expectancy (D) based on a 1-9 Likert scale.

#### CS valence, arousal, and fear ratings

Post habituation phase, no significant differences in valence, arousal, and fear ratings between CS+ and CS- were present, and this was consistent across bromocriptine, placebo and haloperidol groups. Following fear acquisition training, CS+ received significantly higher unpleasantness ratings, higher arousal ratings, and was perceived as inducing more fear than CS- in all three groups (*Figure S3;* *Table S8*). These distinctions between CSs persisted in every group during extinction and recall phases. Post acquisition phase ratings consistently highlighted significant differences between CS+ and CS- in valence, arousal and fear ratings in all groups. Non-parametric ANOVA-type statistic revealed a significant main effect of Stimulus and Phase (main effects: all p<0.001), and a Stimulus × Phase interaction for arousal (F_(2.66,176)=_39.44, p<0.001), valence (F_(2.63,168)=_80.49, p<0.001) and fear (F_(2.63,166)=_41.18, p<0.001) ratings independently of the drug group, as well as a Stimulus x Group x Phase interaction in fear ratings (F_(4.92,166)_=2.66, p=0.021; Table S9). Post-hoc tests showed significant differences between stimuli post fear acquisition training, extinction training and recall phases (least square means tests, Valence: all p<0.001; Arousal: all p<0.001; Fear: all p<0.001), but not following the habituation phase (least square means test, all p>0.99).

#### US unpleasantness, CS/US contingency and US expectancy

Across all participants of the three groups, the likelihood that a US occurred after CS+ presentation was estimated to be 96.40 ± 12.80% (100% probability for 67 out of 75 participants), while after CS- presentation it was 3.87 ± 15.24% (0% probability for 72 out of 75 participants; *Figure S3; Table S5*). Overall median US unpleasantness was rated 7 (IQR 7-8) following acquisition, participants reported recognizing the association between CS+ and US after experiencing an average of 2.27 ± 1.42 electric shocks. No significant differences between groups for both US unpleasantness and CS/US contingency ratings have been found. After the habituation phase, reported US expectancy following CS+ and CS- were not significantly different from each other. Post fear acquisition training, participants reported a higher US expectancy after CS + compared to the CS- and this difference remained until the end of recall (*Figure S3; Table* S8). Non-parametric ANOVA-type statistic revealed a significant main effect of Stimulus and Phase (main effects: all p< 0.001), and a Stimulus × Phase interaction for US expectancy (F_(2.74,195)=_120.13, p<0.001) ratings independently of the drug group (*Table S9*). Post-hoc tests showed significant differences between stimuli post fear acquisition training, extinction training, and recall phase (least square means tests, US expectancy: all p<0.001), but not following habituation (least square means test, p>0.98).

**Table S8.** Self-reported fear evaluation questionnaire ratings in the groups receiving bromocriptine (Brom.), placebo (Plac.) or haloperidol (Halo.) post habituation, fear acquisition training, extinction training, and recall. Median (interquartile range) ratings of valence (A), arousal (B), fear (C) and US expectancy (D) based on a 1-9 Likert scale. Statistically significant differences between CS+ and CS- are shown in bold (least square means tests, p<0.01). Note that no statistically significant differences were found between drug groups and placebo group.

| Stimulus | Time of assessment |  |  |  |  |  |  |  |  |  |  |  |
| --- | --- | --- | --- | --- | --- | --- | --- | --- | --- | --- | --- | --- |
|  | Post Habituation |  |  | Post acquisition |  |  | Post extinction |  |  | Post recall |  |  |
|  | Brom. | Plac. | Halo. | Brom. | Plac. | Halo. | Brom. | Plac. | Halo. | Brom. | Plac. | Halo. |

| *Valence ratings (1 – comfortable, 9 – uncomfortable)* |  |  |  |  |  |  |  |  |  |  |  |  |
| --- | --- | --- | --- | --- | --- | --- | --- | --- | --- | --- | --- | --- |
| CS+ | 5<br>(3-5) | 4<br>(3-5) | 3<br>(3-5) | 7<br>(6-8) | 7<br>(5-7) | 7<br>(6-7) | 4<br>(3-5) | 5<br>(3-5) | 5<br>(2-5) | 4<br>(3-5) | 5<br>(2-5) | 4<br>(2-5) |
| CS- | 5 (3-5) | 5<br>(3-5) | 3<br>(2-5) | 2<br>(1-3) | 3<br>(2-5) | 2<br>(1-3) | 3<br>(2-5) | 3<br>(2-5) | 2<br>(1-4) | 3<br>(2-5) | 3<br>(1-4) | 1<br>(1-3) |
| *Arousal ratings (1 – very calm, 9 – very nervous)* |  |  |  |  |  |  |  |  |  |  |  |  |
| CS+ | 1 (1-3) | 3<br>(2-5) | 2<br>(1-5) | 7<br>(5-8) | 6<br>(5-7) | 6<br>(6-7) | 2<br>(1-4) | 3<br>(1-4) | 3<br>(2-6) | 2<br>(1-5) | 3<br>(1-4) | 2<br>(1-4) |
| CS- | 2 (1-3) | 3<br>(1-5) | 1 (1-3) | 1<br>(1-3) | 2<br>(1-3) | 1<br>(1-2) | 1<br>(1-2) | 1<br>(1-2) | 1<br>(1-2) | 1<br>(1-1) | 1<br>(1-2) | 1<br>(1-1) |
| *Fear ratings (1 – not afraid, 9 – very afraid)* |  |  |  |  |  |  |  |  |  |  |  |  |
| CS+ | 1<br>(1-1) | 1<br>(1-4) | 1<br>(1-1) | 6<br>(4-7) | 4<br>(2-6) | 6<br>(4-6) | 2<br>(1-3) | 2<br>(1-3) | 2<br>(1-5) | 2<br>(1-3) | 2<br>(1-2) | 2<br>(1-5) |
| CS- | 1<br>(1-1) | 1<br>(1-2) | 1<br>(1-1) | 1<br>(1-2) | 1<br>(1-2) | 1<br>(1-1) | 1<br>(1-2) | 1<br>(1-1) | 1<br>(1-1) | 1<br>(1-1) | 1<br>(1-1) | 1<br>(1-1) |
| *US expectancy ratings (1 – US not expected, 9 – US surely expected)* |  |  |  |  |  |  |  |  |  |  |  |  |
| CS+ | 1<br>(1-2) | 1<br>(1-1) | 1<br>(1-1) | 9<br>(9-9) | 9<br>(9-9) | 9<br>(9-9) | 2<br>(1-5) | 3<br>(1-5) | 3<br>(1-5) | 2<br>(1-4) | 2<br>(1-5) | 2<br>(1-5) |
| CS- | 1<br>(1-1) | 1<br>(1-1) | 1<br>(1-1) | 1<br>(1-2) | 1<br>(1-2) | 1<br>(1-1) | 1<br>(1-2) | 1<br>(1-2) | 1<br>(1-1) | 1<br>(1-2) | 1<br>(1-2) | 1<br>(1-1) |

**Table S9.**
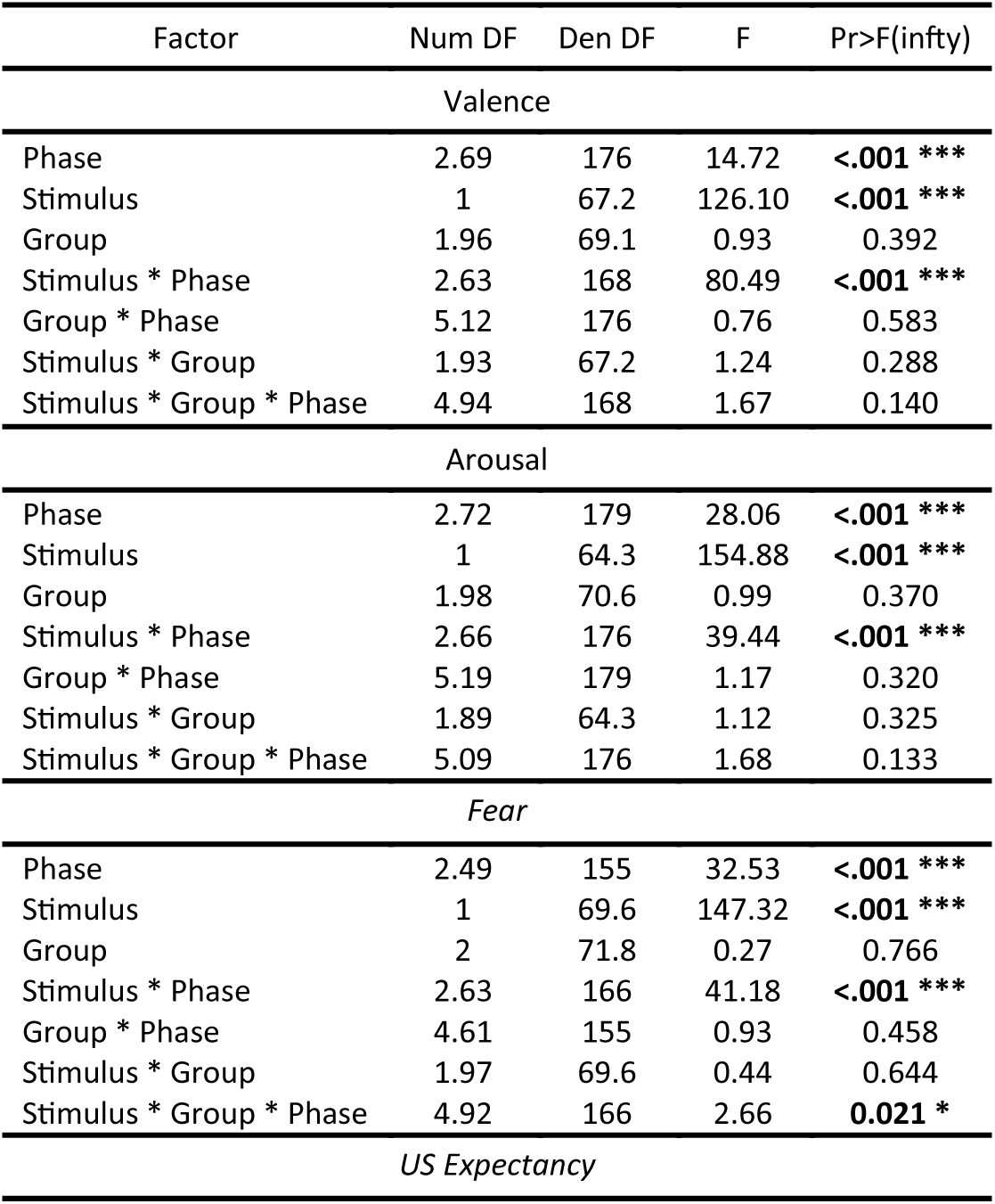
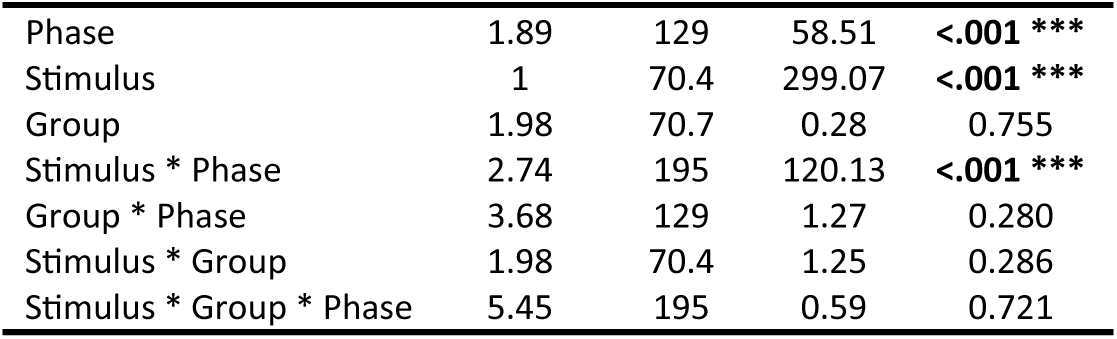
Self-reported fear evaluation questionnaire ratings in the groups receiving bromocriptine, placebo or haloperidol. Results of the non-parametric ATS for repeated measures on all phases. (*p<0.01; ** p<0.05; *** p<0.001)

### Statistics for pupillometry and skin conductance responses

**Table S10.**
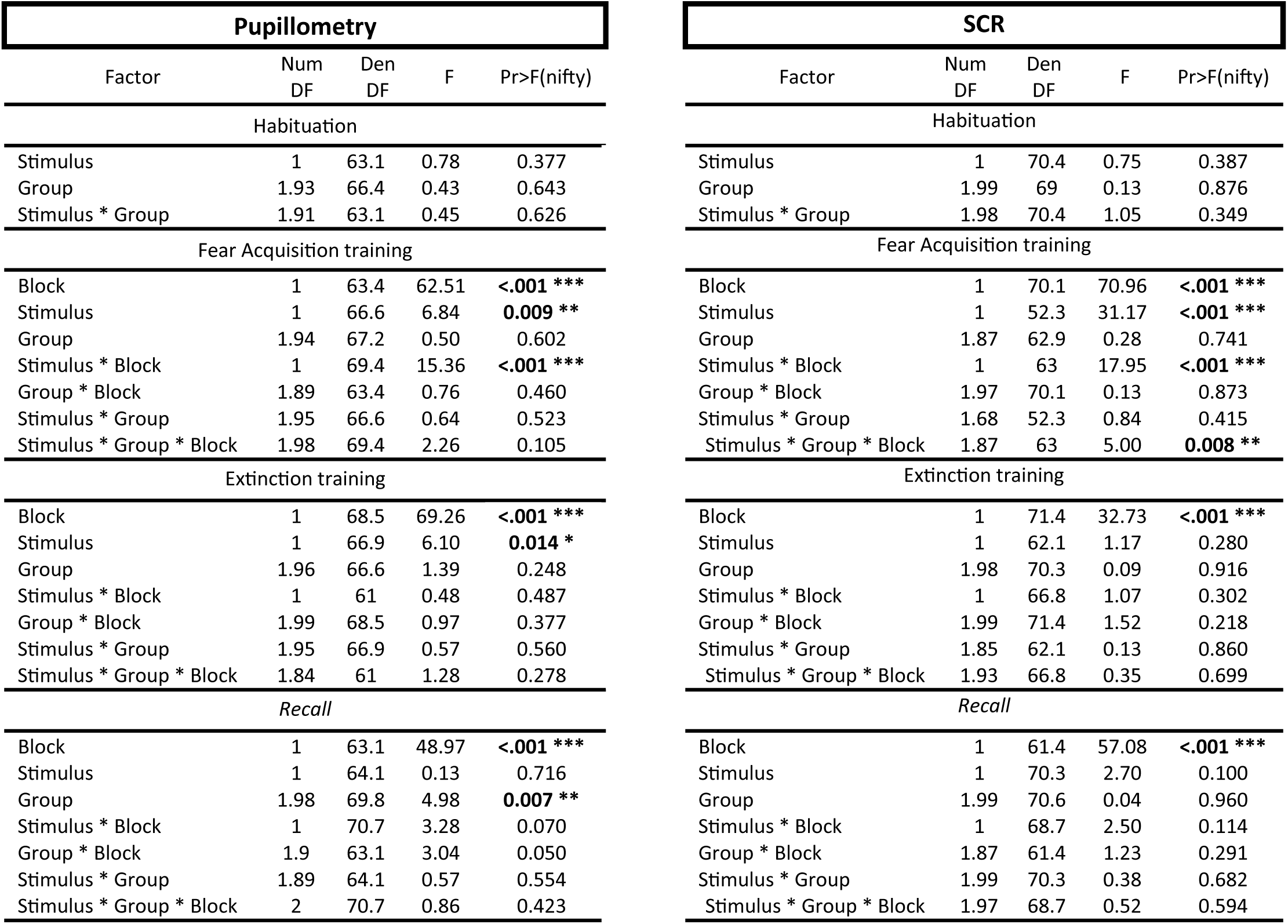
Pupil size and SCR statistics in the groups receiving bromocriptine, placebo or haloperidol. Results of the non-parametric ATS for repeated measures on all phases. (*p<0.01; ** p<0.05; *** p<0.001)

### First recall trials pupillometry and skin conductance responses

**Figure S4.**
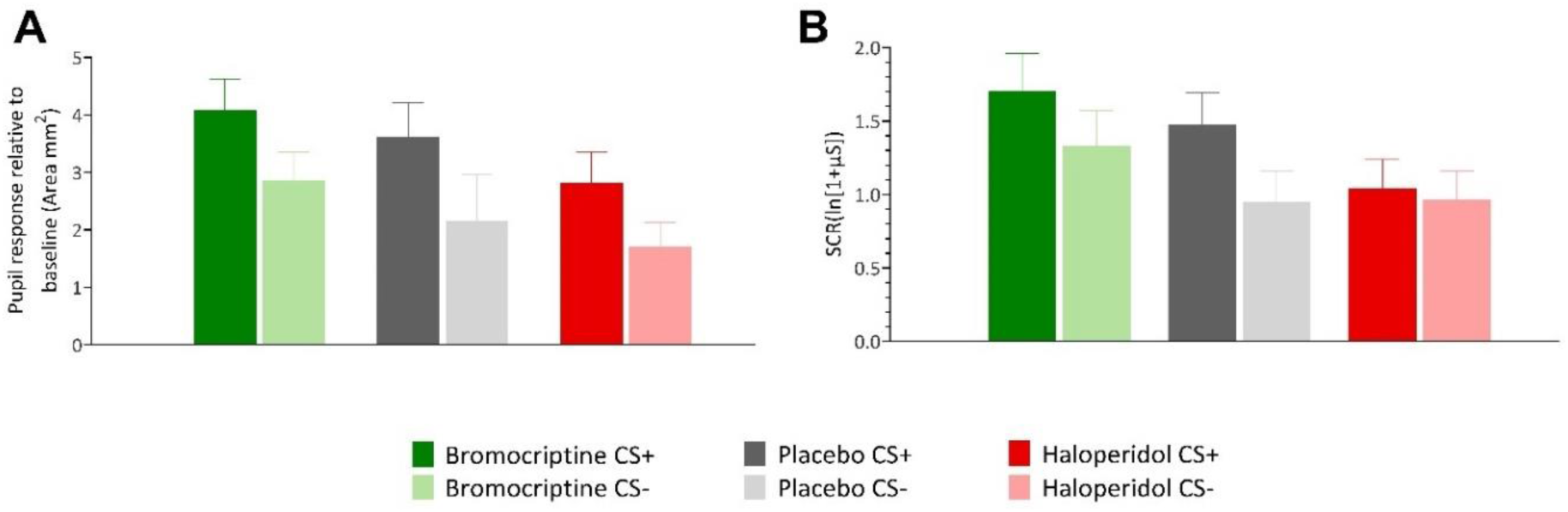
(A) Pupil responses relative to baseline for first recall trial in the bromocriptine, placebo and haloperidol groups. Bars represent means, error bars indicate S.E.M. (B) Skin conductance responses (SCRs) for first recall trial in the groups receiving bromocriptine, placebo and haloperidol. Bars represent means, error bars indicate S.E.M.

We assessed the first trial in recall across the bromocriptine, placebo, and haloperidol groups using both pupil dilation and SCR (*Figure S4–A and B; Table S11*). Both measures showed a significant difference between CS+ and CS- trials (Pupil: F_(1, 54.6)_=8.23, p=0.006; SCR: F_(1, 68.5)_=8.66, p=0.003), indicating spontaneous recovery of the initial fear association. However, there were no significant differences between the drug groups for either measure (Pupil: F_(1.93, 62.1)_=2.96, p=0.054; SCR: F_(1.99, 69.8)_=1.42, p=0.243). For the interaction between stimuli and drug group, neither measure showed a significant effect (Pupil: F_(1.77, 54.6)_=0.13, p=0.857; SCR: F_(1.97, 68.5)_=1.37, p=0.255;*Table S11*).

**Table S11.** Pupillometry and SCR statistics in the groups receiving bromocriptine, placebo and haloperidol. Results of the non-parametric ATS for repeated measures on the first trial of recall. (*p < 0.01; ** p < 0.05; *** p < 0.001)

| Factor | Num DF | Den DF | F | Pr>F(infty) |
| --- | --- | --- | --- | --- |
| **Pupillometry** |  |  |  |  |
| Stimulus | 1 | 54.6 | 8.23 | **0.004\*\*** |
| Group | 1.93 | 62.1 | 2.96 | 0.054 |
| Stimulus * Group | 1.77 | 54.6 | 0.13 | 0.857 |
| **SCR** |  |  |  |  |
| Stimulus | 1 | 68.5 | 8.66 | **0.003\*\*** |
| Group | 1.99 | 69.8 | 1.42 | 0.243 |
| Stimulus * Group | 1.97 | 68.5 | 1.37 | 0.255 |

### Pupil size variation at baseline

**Table S12.**
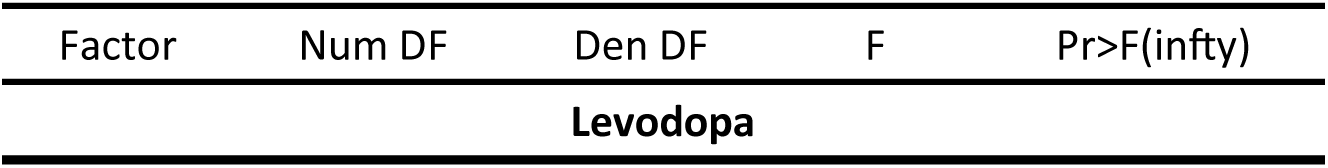
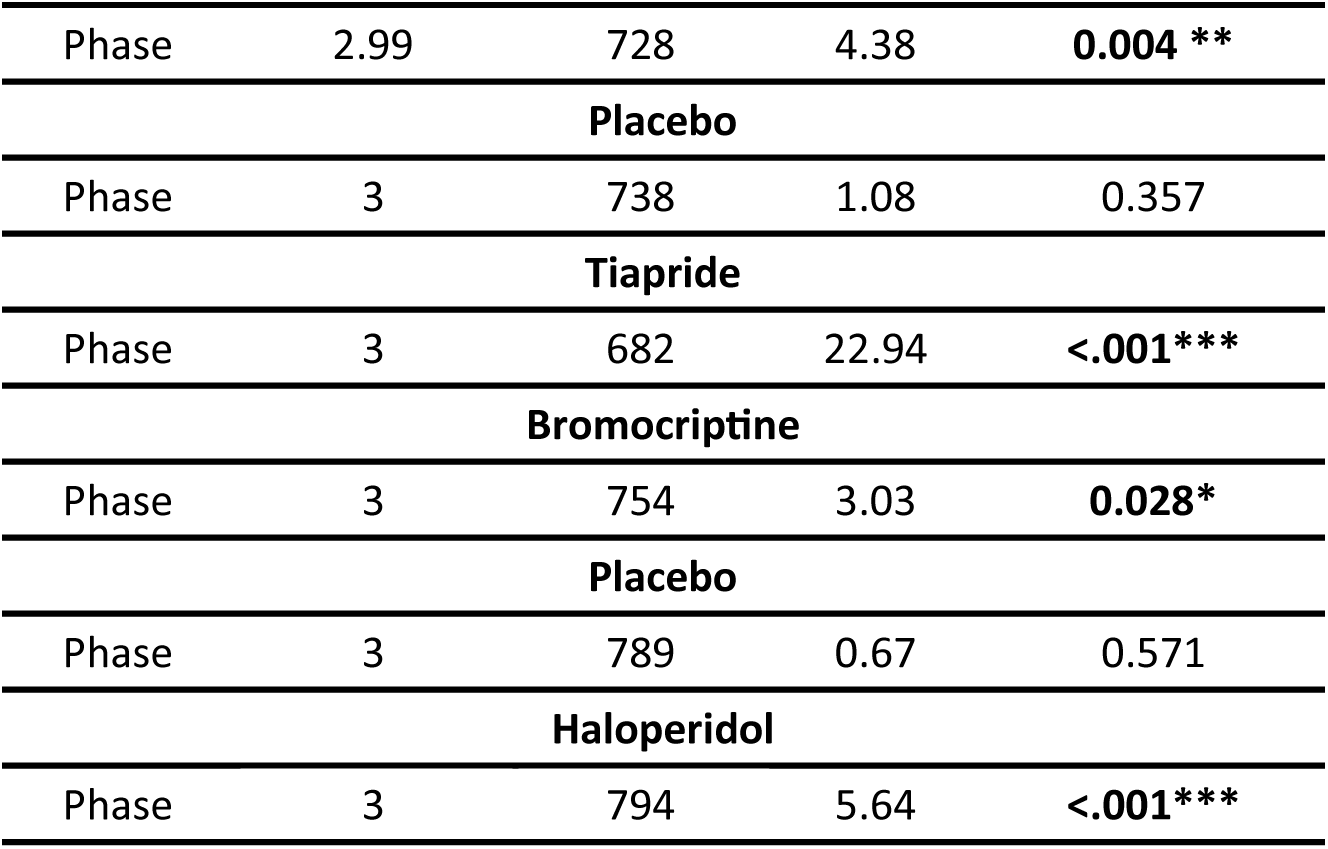
Pupil size variation (in mm^2^) 2s prior CSs onset during the first 8 trials in both group A receiving levodopa, placebo or tiapride and group B receiving bromocriptine, placebo or haloperidol. Results of the non-parametric ATS for repeated measures between all phases inside each group. (*p<0.01; ** p<0.05; *** p<0.001)

