## Supplementary figures and images for "Dopaminergic drugs modulate fear extinction related processes in humans, but effects are mild"

### Figure 2

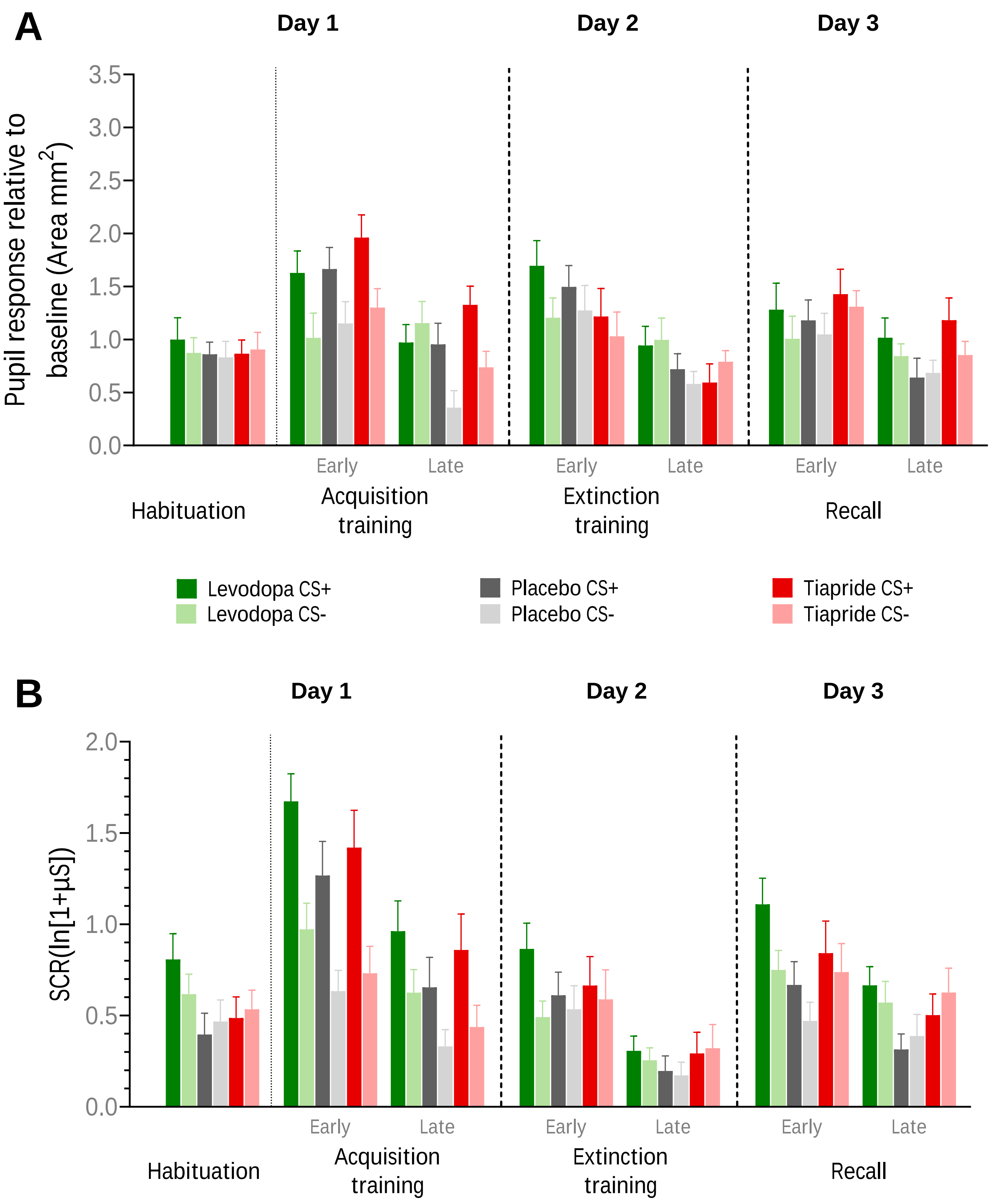

### Figure 3

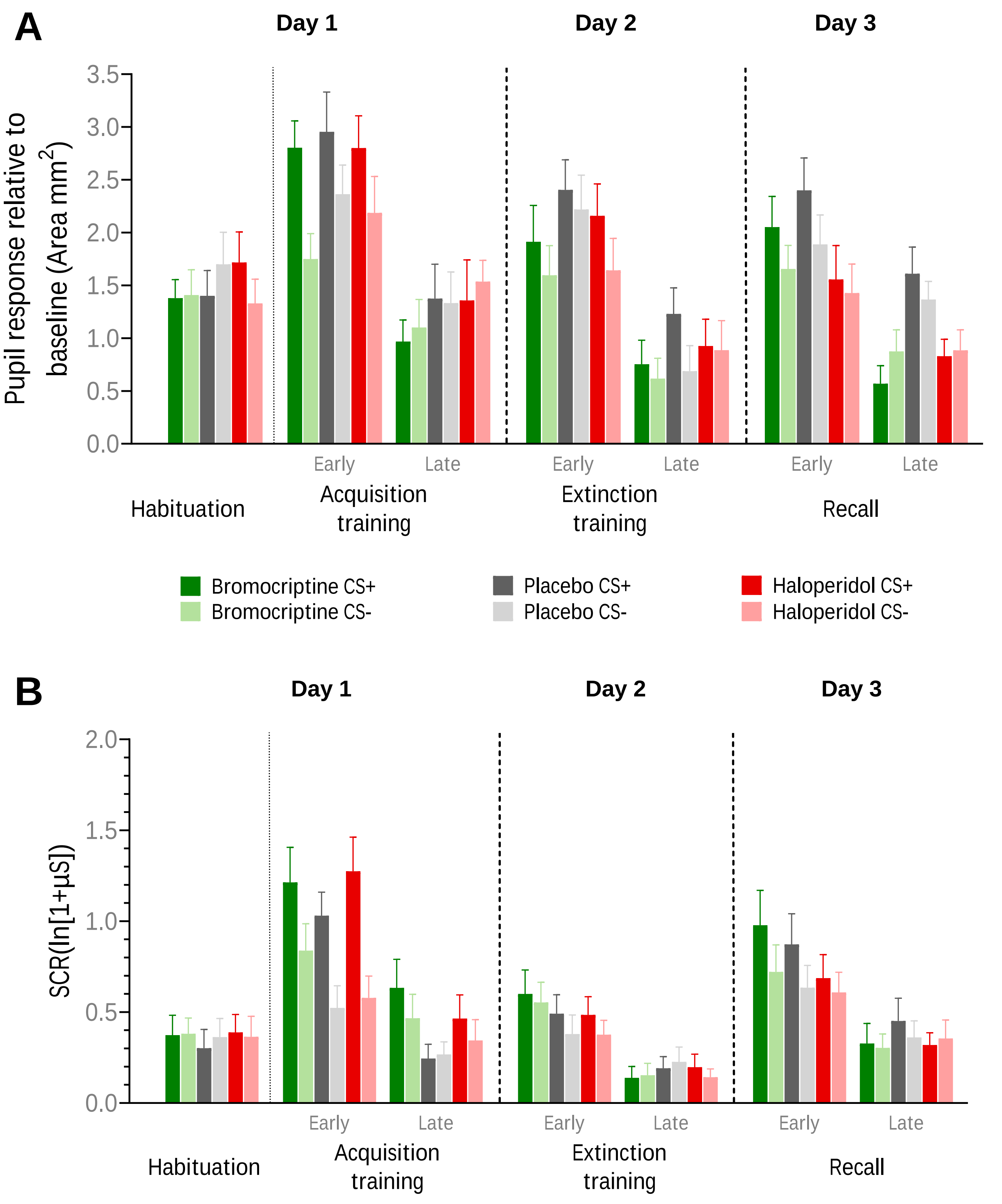

### Figure 4

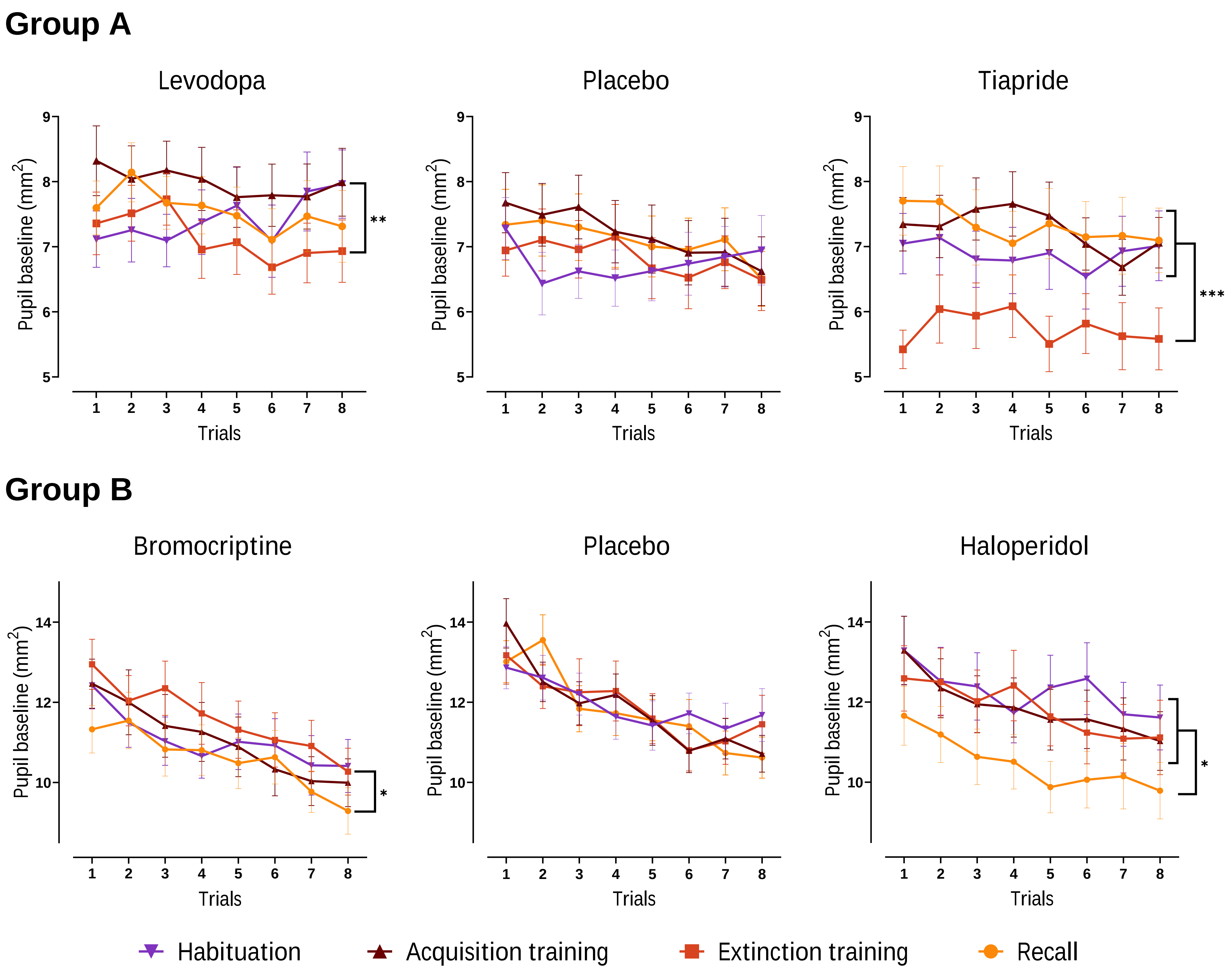

### Supplementary Figure S1

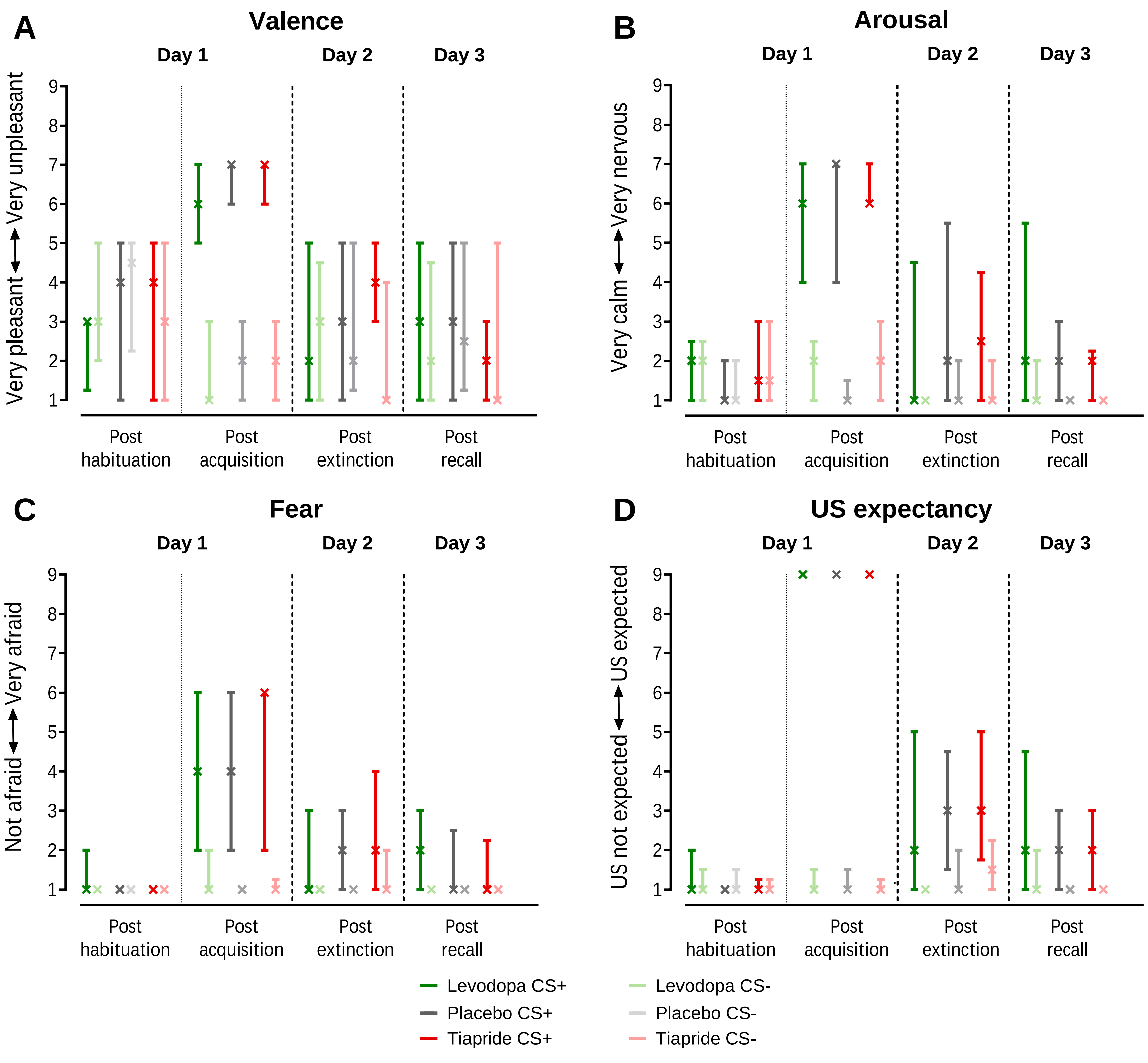

### Supplementary Figure S3

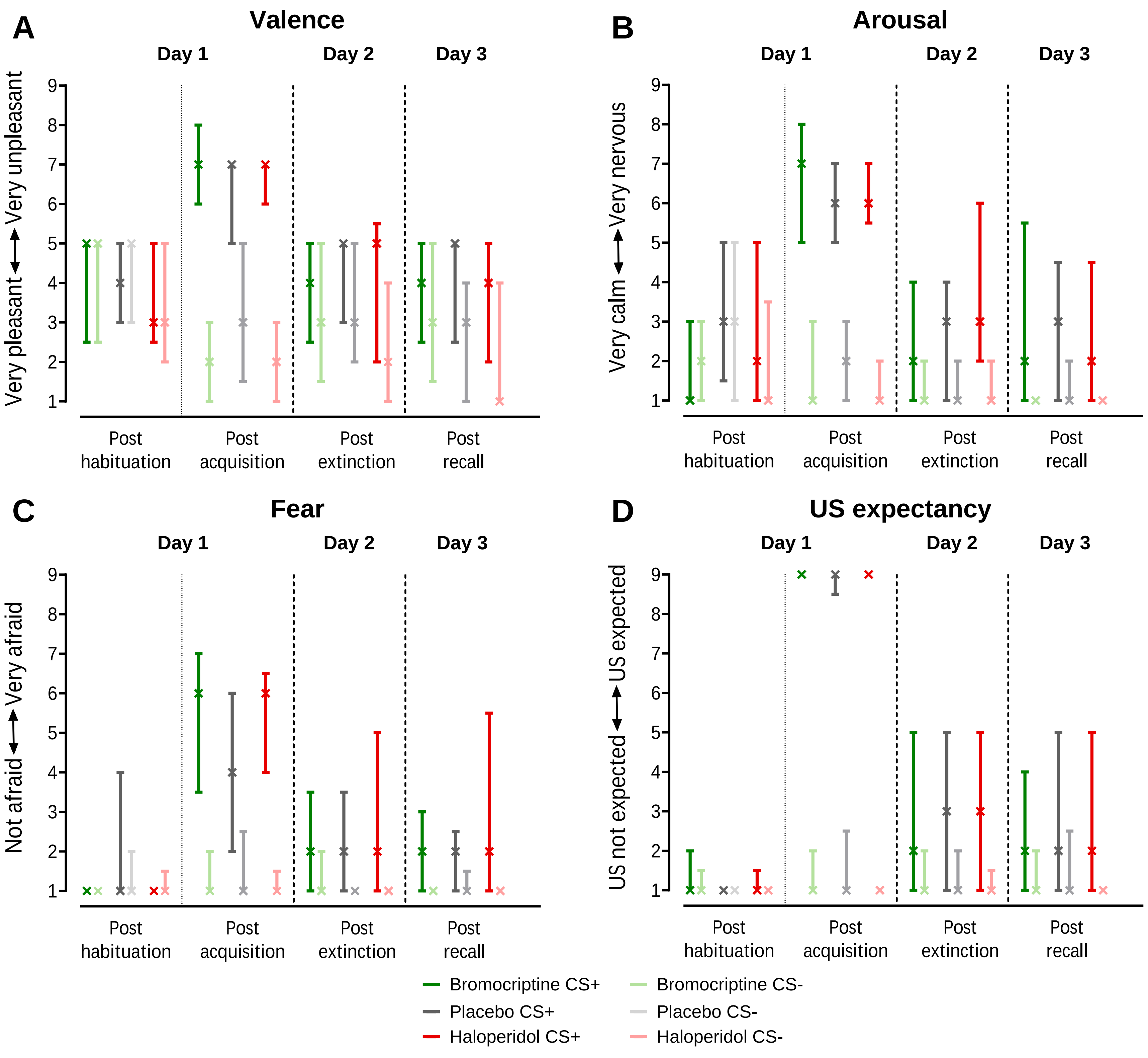
